# Targeting Nitric Oxide Synthase 2 Reverses Learning Deficits in an Oligodendrocyte-Focused Model of Costello Syndrome

**DOI:** 10.64898/2026.06.01.729333

**Authors:** Celeste Martinez, Saul Lopez, Daniella Hernandez, Ashley Figueroa, Karla D. López-Lorenzo, Jonathan Rodriguez, Vijay Radhakrishnan, Montserrat Rodríguez-Rodríguez, Anderson M. Winkler, Abraham Cisneros-Mejorado, Akiko Nishiyama, Alejandro López-Juárez

## Abstract

Oligodendrocyte (OL) functions greatly improve information processing in the brain; consequently, compromised OL biology negatively impacts brain function in various diseases. Young-adult Costello Syndrome (CS) patients present with learning disabilities and regression of psychomotor skills in correlation with abnormal brain white matter, delayed myelination, and dysmyelination. This suggests the involvement of OLs and myelin in abnormal brain physiology during and beyond development; however, the neuropathophysiology of CS remains poorly understood. Replacing the endogenous *HRas* gene with the CS-causing *HRasG12V* mutant gene in adult OLs (pHRasG/+ mice) induces structural myelin abnormalities; nonetheless, any functional impact is unknown. Here we show robust learning - but not memory - deficits in pHRasG/+ males and modest, delayed learning issues in females. Learning phenotypes are transient, likely involving compensatory responses from the OL lineage and microglia. Diffusion-weighted magnetic resonance imaging revealed regional time- and sex-dependent compromised myelin microstructure, coinciding with the peak of learning issues. Remarkably, transcripts associated with inflammation/immune function are upregulated and pharmacological control of nitric oxide synthase 2 restores learning in male pHRasG+/- mice. Our study supports the notion that abnormal nitric oxide signaling in mature OLs affects learning, suggests a chronic impact of glia-myelin mechanisms in CS neuropathology, and proposes molecular targets with therapeutic potential.

## 1. INTRODUCTION

Mutations in genes involved in the RAS/MAPK pathway cause RASopathies, a group of diseases that affect 1/1000 individuals worldwide ^1^. Among the diverse body systems affected, RASopathy patients present with compromised brain function ^2^. The RASopathy Costello Syndrome (CS) is caused by hyperactivating mutations in a single allele of the *HRas* gene ^3^, and patients present with various degrees of learning disabilities, delays or regression in psychomotor skills ^4–7^, and predisposition to ASD and ADHD ^8–11^. In parallel, CS patients show brain white matter (WM) abnormalities including small corpus callosum (CC) ^12^, hypodense WM ^13,14^, delayed myelination ^15,16^ or dysmyelination ^14,17^. Of note, psychomotor regression and dysmyelination findings in CS suggest pathological mechanisms continuing beyond development. Age and sex may influence specific areas of cognition in CS, adding complexity to its mechanisms ^18^. While brain cortex development seems to be affected by CS mutations in research models ^19^, developmental WM/myelin abnormalities and their repercussions in adult patients or models remain poorly explored.

In the brain, compact myelin produced by oligodendrocytes (OLs) wraps around neuronal axons, speeding up and synchronizing action potentials. This apparently simple scheme increases the speed of conduction in the brain in >100 fold ^20^, mainly through WM tracts that can occupy over half of the brain volume in humans ^21^. Beyond development, changes in adult OLs, myelin, and WM continue and impact brain function in health and disease ^22–24^. For example, learning challenges in rodents increase OL numbers, and blocking OL differentiation prevents learning ^25–28^; hence, it was proposed that new myelin underlies WM structural changes that follow learning in humans ^28–30^. Furthermore, roles of OLs in regulating brain function are proposed through their metabolic support to axons ^31,32^ and apparent immune functions ^33,34^. In parallel, it was proposed that defective OL biology underlies neurological conditions including learning disability, attention deficit and hyperactivity disorder, autism spectrum disorders, etc. ^35–37^. As CS patients present with learning issues and abnormal WM/myelin, inducing CS-causing mutations in mice represents a relevant strategy to uncover the involvement of OLs in impaired learning. Various paradigms ^38–40^ including the fine motor skill learning complex wheel (CW; running wheel with unevenly spaced rungs) test, have been used to show that OLs regulates learning ^26,27,41^. The CW voluntary nocturnal test ^42^ relies on the integrity of the CC ^43,44^ as mice adopt strategies for asymmetrical gait running on wheels with unequally distributed gaps ^26^. CW learning seems to involve additional circuits beyond those required for symmetric gait dependent on hippocampal integrity ^26,45^. Overall, mice running on CWs improve their skills every night and eventually run as fast as in regular wheels ^27^. Therefore, we tested whether abnormal OL biology triggered by a CS-causing mutation impairs CW learning.

The *HRasG12V* gain of function mutation strongly activates the RAS pathway ^46,47^ and drives severe CS presentation ^3,48^. We reported that replacement of the normal *HRas* with *HRasG12V* in its endogenous locus (thus preserving genomic environmental regulation) in myelinating Plp1+ cells (pHRsG/+ mice), hyperactivates the RAS/MAPK, nitic oxide (NO), and Notch pathways, and impacts myelin g-ratio and compaction ^49^. Here we used pHRsG/+ mice to test whether compromised OL biology alters brain cell physiology and function. We found acute CW learning defects in mutant males, and less severe issues in females; in both sexes, learning phenotypes were transient likely due to compensatory mechanisms involving the OL lineage and microglia. We revealed NO synthase 2 (NOS2) activity as a key mediator of learning issues. Moreover, diffusion-weighted magnetic resonance imaging (dMRI) showed transient white matter (WM) microstructural defects in pHRsG/+ males, synchronized with the period of learning impairment. Overall, our results establish temporal associations for abnormal OL biology and learning defects that can be precluded by NOS2 inhibition. Whether such associations are chronic in the context of germline *HRas* mutations remains to be investigated.

## 2. METHODS

### 2.1. Animals

All studies in mice were approved by the University of Texas Rio Grande Valley Institutional Animal Care and Use Review Committee (IACUC). Mouse strains were validated and alleles selected upon genotyping as previously described for FR-HRasG12V ^50^, PlpCreERT2 ^51^, and cag-catEGFP ^52^ alleles. The mouse genotypes used were *PlpCreERT2*;*FR-HRasG12V/+* (pHRsG/+) and *PlpCreERT2* or *FR-HRasG12V/+* which were phenotypically wild type (WT) and indistinguishable from each other. The CreERT2-inducible GFP reporter gene cag-catEGFP was included in pHRsG/+ or WTs alleles to study the fate of recombinant cells. All strains were back crossed for >10 generations with the C57BL/6J strain background. Mice were housed at a controlled temperature and humidity on a 12 h light–dark cycle and with free access to food and water. Body weight (BW) was measured in all individuals before tamoxifen treatment, the first day running in CWs, as well as day 36 (reintroduction to CWs), and day 42 (end of CW test); mice with variations in BW (>10%) that suggested compromised health were not included in experimental groups.

### 2.2. Drug Treatments

All treatments were equally administered to both pHRsG/+ and WT control mice. For tamoxifen (tmx)-mediated recombination/mutation of *HRas* and concomitant GFP expression, 2-month-old (2MO) mice were injected intraperitoneally with tmx (75 mg/kg of body weight, in sunflower seed oil; Sigma-Aldrich, St. Louis, MI, USA) twice daily, for 3 consecutive days. For 1400w (N-([3-(aminomethyl)phenyl]methyl)ethanimidamide) treatment, the drug was acquired from MedChemExpress (NJ, U.S.) and diluted in 10% DMSO/90% Corn Oil (Sigma-Aldrich, St. Louis, MI, USA) at 0.08mg/ml; a dose of 0.2mg/kg of body weight was subcutaneously administered to mice at the schedules detailed in results. Pexidartinib (PLX3397; Chemgood, VA, USA) was prepared in the Rodent Diet AIN-76A (290 mg/kg diet, Research Diets Inc., NJ, USA) and administered ad libitum at the schedules detailed in the results.

### 2.3. Complex Wheel Test

Individual mice of the appropriate age, genotype, and treatment were introduced to single cages with a CW (wheels with 22 unevenly spaced rungs) at the beginning of the 12-hour dark cycle, and running was monitored 24/7 with an automated infrared system (SAS: Lafayette Instruments, IN, USA). The first CW introduction was for 2 weeks, then mice were housed without wheel for 3 weeks, and reintroduced to the CWs for 1 week to evaluate memory of skills previously acquired. Researcher influence was minimized, with cages opened only for food/water replenishment. The running bin was in meters/minute and analyzed per 12-hour dark period (or nights; Ns) for the duration of the experiment (overall 42 nights). The total average speed (TAS) comprises all minutes per night, while the activity average speed (AAS) accounts only for minutes in which mice run > 1 meter. Maximum Speed (MaxSpd) is the average of the top three speeds reached in a single night, and the activity reports the number of minutes running at least one meter. Nights in the 1^st^ introduction (N1-14) and 2^nd^ introduction (N36-42) were independently analyzed. Light intensity was homogeneous for all cages, and cage location was randomized. Whiskers were trimmed before N1 and N36 to avoid tactile anticipation of missing rungs.

### 2.4. Statistics

Raw CW data obtained with the SAS software (version 23; Lafayette Instruments, IN, USA) or from CC image quantifications were processed by investigators blinded to the mouse genotype/treatment. All quantifications are presented as mean +/− standard error of the mean. Data were analyzed by two-way ANOVA with gender, genotype, or time as sources of variation. For two-group comparisons (single night in CWs analyses and for pHRsG/+ vs. WT image evaluations) the Bonferroni’s post hoc (after two-way ANOVA) test and/or unpaired Student’s t-tests were used. All statistical analyses were performed using Prism 9 software (GraphPad, Boston, MA). Significant differences were considered at P < 0.05 (*). For CW analyses, four parameters were obtained for each 12h dark period. The CW average speed of phenotypically WT mice had an approximately normal distribution (P = 0.2, K-S test) that, together with the low data variability, shows significant differences using a low “n” of mice ^26,27^; to account for the potential impact of phenotype variability caused by the *HRas* mutation, a minimum “n” of 5 mice/gender/genotype was established.

### 2.5. Brain Dissection, Immunostaining, and Confocal Imaging

Mice were overdosed with isoflurane and perfused with 1X PBS followed by 4% paraformaldehyde. Brains were dissected, post-fixed in 4% paraformaldehyde for one night, and sectioned (30μm) using a vibratome (Leica, Wetzlar, Germany). Floating sections were processed for immunodetection using antibodies for the reporter gene GFP (Nacalai Tesque, Kyoto, Japan), and the markers GSTpi (MBL Ltd, Tokio, Japan), CC1 (Calbiochem, San Diego, CA, USA), NG2 (Millipore, Burlington, MA, USA), PDGFRa (Rn’D systems, Minneapolis, MN), GFAP (Dako, Santa Clara, CA, USA), IBA1 (Wako, Osaka, Japan), NeuN (Millipore, St. Louis, MO), and pERK (Cell signaling, MA, USA). Fluorophore-conjugated secondary antibodies (Cy2, Cy3, and Cy5; Jackson ImmunoResearch, West Grove, PA, USA) were used to detect the primary antibodies using a Leica 2500SPE (Leica Microsystems, Wetzlar, Germany) or a Nikon AXR (Nikon Instruments, NY, USA) confocal microscopes. Immunodetected cells were automatically (nuclei, GFP, pERK, GSTpi, and CC1) or manually (IBA1, GFAP, and PDGFR) quantified from seven regions (**Fig. 1C**) throughout the anterior-posterior (I-IV) and central “C”, lateral “B” axes of the corpus callosum (CC), using the software ImageJ-ITCN plugin (NIH, MD, USA); data was the average from all regions unless otherwise disclosed. For automatic counting, a region of interest (ROI) of ∼1mm^2^ (Arbitrary Unit; AU) was established for all images and the total number of marker+ cells was automatically calculated using equal ImageJ/ITCN parameters for size, distance between cells, and signal threshold per marker. The number of positive cells was normalized to the total number of Dapi+ nuclei in each ROI, to minimize the effect of region variability in cell density.

**Figure 1.**
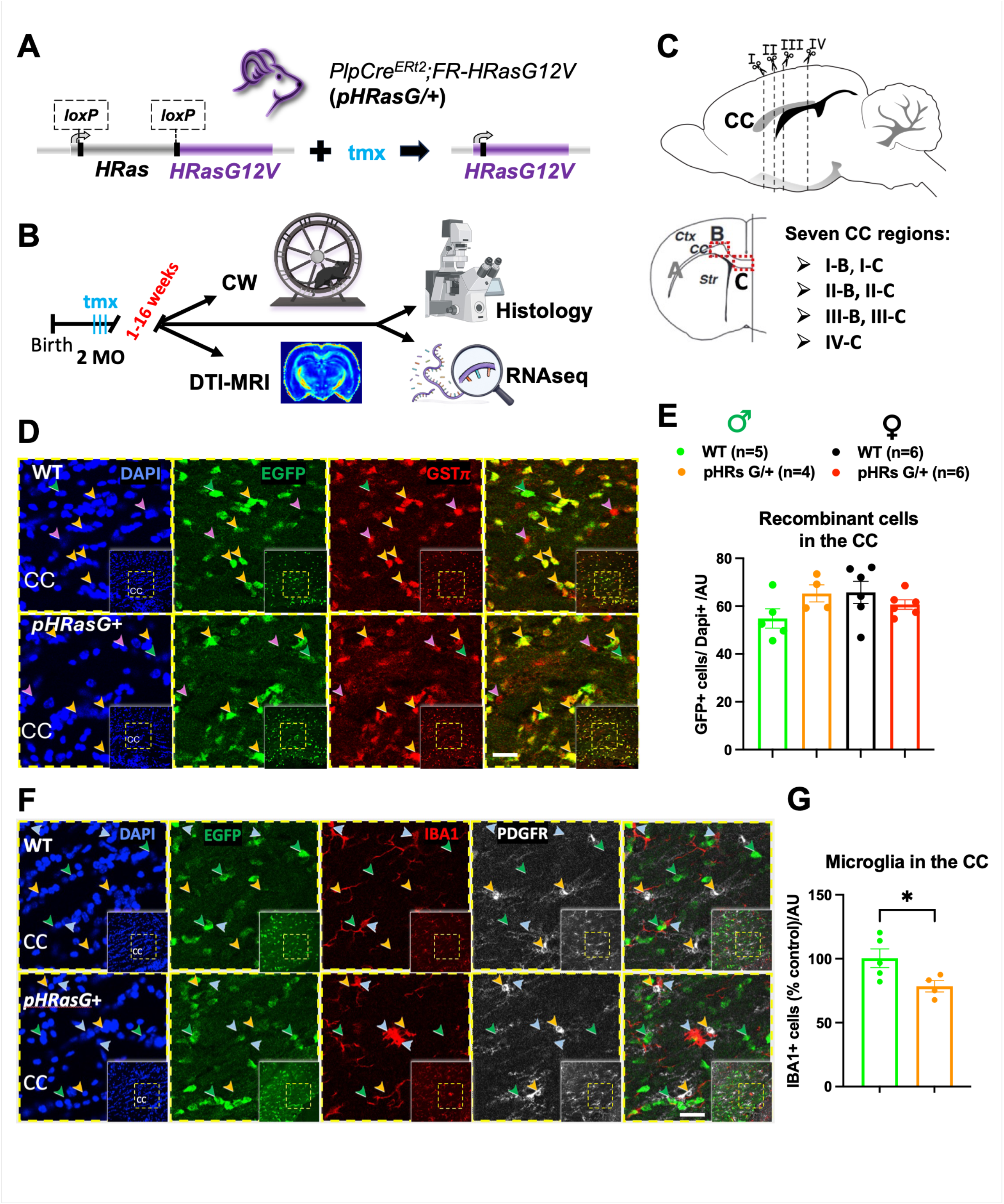
Specific *HRas* Mutation in OLs Triggers Non-Cell-Autonomous Responses. **A)** Recombination strategy for tamoxifen (tmx; cyan)–mediated replacement of the *HRas* gene (grey, flanked by loxP sites) with *HRasG12V* (purple) in *PlpCreERt2;FR-HRasG12V* (pHRsG/+) mice. **B)** 2-month-old (2MO) mice treated with tmx are subjected to the complex wheel (CW) test, histological, bulk RNAseq, and DTI-MRI analyses, 1-16 weeks later. **C)** Cell populations are analyzed in seven regions throughout the anterior-posterior (I-IV) and central “C”-lateral “B” axes of the corpus callosum (CC), unless otherwise disclosed. **D)** Immunostaining of coronal sections showing recombinant cells (GFP+, green arrowheads), OLs (GSTpi+, purple arrowheads), and GFP+GSTpi+ recombinant OLs (orange arrowheads) in the CC of WT and pHRsG/+ mice. **E)** The percentage of recombinant cells (normalized to dapi) in WT and pHRsG/+ mice is not significantly different (unpaired Student’s *t* test; males P= 0.14 and females P= 0.24). Genotype/sex color code and “n” per group in **E, G**: male WT (green) and pHRsG/+ (orange); female WT (black) and pHRsG/+ (red). **F)** Representative immunostaining indicates recombinant cells (green arrowheads), microglia (IBA1+, blue arrowheads), and OPCs (PDGFRa+, orange arrowheads) in the CC of WT and pHRsG/+ mice. **G)** The percentage of IBA+ cells in WT and pHRsG/+ mice (% of age/gender matched WTs) indicates significantly decreased # of microglia in male pHRsG/+ mice (unpaired Student’s *t* test; P= 0.047). Insets in D and F show the CC region in the high magnification image (dotted yellow square). Dapi was used to stain nuclei and normalize cell densities per arbitrary units (AU). Scale bar = 25μm. *p < 0.05.

### 2.6. Bulk mRNAseq

Optic nerves from WT and pHRasG/+ mice (n=4/genotype) were collected 2w post-tmx, total RNA was extracted using ZR RNA Microprep (Zymo cat # R1060) and submitted to Novogene Inc. (Sacramento, CA, U.S.A.) for library preparation and sequencing. The RNA concentration and integrity was assessed by NanoDrop spectrophotometry and Agilent 2100 Bioanalyze (RIN>8.5). mRNA was enriched, fragmented, and reverse-transcribed using random hexamer primers. Following second-strand synthesis, libraries were prepared (end repair, A-tailing, adapter ligation, size selection, amplification, and purification), quantified (Qubit fluorometry and rtPCR), validated (Agilent Bioanalyzer), and sequenced (Illumina NovaSeq X Plus platform; paired-end 150-bp reads). Raw FASTQ files and downstream analyses were processed using Novomagic (Novogene, CA, US). Low-quality reads were removed. Sequencing quality was evaluated using error-rate, GC-content, Q20, and Q30 metrics. Reads were aligned (GRCm39/mm39; HISAT2) and used for gene quantification. Gene expression was quantified as raw read counts and FPKM values. Samples were evaluated by Pearson correlation and principal component analysis. Differential expression analysis was performed with DESeq2 using normalized read counts and negative binomial model. Statistical significance was determined by Wald tests with Benjamini–Hochberg false discovery rate correction. Genes with an absolute log_2_ fold change ≥1 and adjusted *P* ≤0.05 were considered differentially expressed. Functional analysis of DEGs included GO and KEGG Enrichment. Sequencing data are publicly available on the GEO database (GSE_).

### 2.7. Diffusion-Weighted Magnetic Resonance Analysis (dMRI)

Diffusion-weighted magnetic resonance imaging and analyses of fixed mouse brains were performed at the Laboratorio Nacional de Imagenología por Resonancia Magnética (LANIREM, QRO, Mex.) using a 7 T magnet 70/16 scanning system (Bruker Pharmascan, MA, US), interfaced to a Paravision 7.0.1 console (Bruker, Ettlingen, Germany), and with a Helium-cooled two-channel rat-head coil (Bruker Cryoprobe, Germany). Data sets were acquired using a spin-echo, single-shot echoplanar imaging sequence with the following parameters: 30 sagittal slices of thickness = 0.5 mm, slice gap = 0.1 mm, repetition time = 2 s, echo time = 20 ms, FOV = 15×30 mm^2^, image size = 128×128 mm. A multi-shell acquisition was used, with 30 diffusion directions and b-values 650 and 1250 s/mm^2^, 1 average. Pre-processing of data sets included both reduction of motion and eddy current-induced geometric distortions by linear transformation of each volume to the average non-diffusion weighted volume and denoising via random matrix theory ^53^. The MRtrix software package (http://www.mrtrix.org) was used to estimate the diffusion tensor model, from which fractional anisotropy (FA) and principal diffusion vector (PDV) maps were derived. These dMRI parameters were analyzed in regions of interest (ROI) manually delineated on PDV images. Four regions, including the anterior (aCC) and posterior (pCC) corpus callosum, in the central and lateral areas, were included in the analysis.

## 3. RESULTS

### 3.1 Specific *HRasG12V* Mutation in OLs Causes Cell-Autonomous and Non-Cell-Autonomous Effects in the Corpus Callosum

To investigate the impact of *HRasG12V* mutation in myelinating cells on brain physiology and function, the *HRas* gene was mutated in adult Plp1-expressing cells and phenotypes were analyzed 1-16 weeks later (**Fig. 1A-B)**. Two-month-old (2MO) mice carrying the ‘‘flox-and-replace’’ *HRasG12V* ^50^ allele (henceforth WT), or both *HRasG12V* and *PlpCreER* ^51^ alleles (henceforth pHRsG/+) were treated with tamoxifen (tmx) to induce *HRas* mutation; *HRasG12V* is only expressed when tmx induces excision of the endogenous wild type *HRas* gene flanked by loxP sites (**Fig. 1A**). To track the fate of recombinant/mutant cells, a subset of mice also carried a tmx-inducible EGFP reporter gene ^52^. One-week (1w) post-tmx treatment, we immuno-detected GFP signals in an average 54.85 +/- 8.98% and 65.31 +/- 7.12% of total cells (Dapi+) across seven regions of the corpus callosum (CC, **Fig. 1C**) of WT and pHRsG/+ male mice, respectively (**Fig. 1D-E**). Similarly, 65.79 +/- 11.29% and 60.69 +/- 4.86% of total cells in the CC of WT and pHRsG/+ females showed GFP signals (**Fig. 1E**). No significant differences in the number of recombinant cells were detected between genotypes or sexes, indicating efficient recombination with no effects of mutation status or sex. Analysis of cells showing staining for both GFP and the OL marker GSTpi, indicated that similar proportions of GSTpi+ OLs were recombinant throughout the CC regions in WT and pHRsG/+ males (63.02% +/- 11.55% and 62.96 +/- 11.54%, respectively). Female WT and pHRsG/+ mice showed similar numbers of GFP+GSTpi+ cells among GSTpi+ OLs (74.29% +/- 7.42% and 70.02% +/- 7.06%, respectively). On the other hand, among GFP+ cells 51.45 +/- 8.23% and 51.67 +/- 10.08% showed signals for GSTpi in WT and pHRsG/+ males, respectively. This apparently low number of recombinant OLs could reflect low GSTpi detection threshold in an OL subpopulation (example in **Fig. 1D**; green arrows). To confirm that PlpCreER recombination was specific to OLs, co-immunostaining for GFP and NeuN+ neurons, GFAP+ astrocytes, PDGFRa+ OPCs, and IBA1+ microglia were performed. Comprehensive confocal microscopy analysis revealed virtually no overlap between GFP+ signals with neuron, astrocyte, OPC (**Sup. Fig. 1A-D**), or microglia (**Fig. 1F**) markers. Interestingly, we observed that microglia morphology and density changed in pHRsG/+ males. The percentage of IBA1+ cells showed a significant reduction (21.53% +/- 8.8%) in the CC as compared with age-matched WTs (**Fig. 1G**), and cells adopted an amoeboid morphology; microglial activation may thus contribute to the apparent microglia number reduction. A regional increase in astrocytes in the lateral CC (**Sup. Fig. 1E**, regions I-B to III-B) and a decrease in OPCs confined to the anterior central CC (**Sup. Fig. 1F**, region I-C) were detected. Overall, we observed efficient and OL-specific recombination 1w post-tmx in WT and pHRsG/+ mice; additionally, widespread non-cell autonomous phenotypes in microglia as well as confined changes in astrocytes and OPCs were present in *HRas* mutant mice. We then explored the impact of *HRasG12V* on learning and memory.

### 3.2 Female pHRsG/+ Mice Show Mild Fine Motor Skill Learning Issues 8-10 Weeks After Mutation

One to six months post-tmx, pHRsG/+ mice do not develop gross brain structural or obvious behavioral issues, and OL numbers as well as myelin wrap counts are normal; however, abnormal myelin g-ratio and compaction were described ^49^. Whether these or other undescribed OL phenotypes impact learning remains unknown, so we used the CW test to evaluate fine-motor skill learning curves in pHRsG/+ mice. Eight to ten weeks (8w-10w) post-tmx treatment, mice were introduced to individual CWs for 2w, then were housed with no wheels for 3w, and introduced to CWs for an additional week (2^nd^ introduction) to evaluate memory of skills learned (**Fig. 2A**). We observed that the nocturnal activity levels (minutes run per 12-hour dark cycle) were overall maintained during the whole experiment in WT and pHRsG/+ males with no significant differences between genotypes (**Sup. Fig. 2A** and **Sup. Fig. 3A-B**). Females of both genotypes showed variable activity, with significantly increased levels starting in the 2^nd^ CW introduction (night 36; N36) as compared with N14 (**Sup. Fig. 2B** and **Sup. Fig. 3C**) and decreased by N42. Importantly, no differences in activity levels were detected between pHRsG/+ and WTs, male or female mice, for either the 1^st^ or 2^nd^ CW introduction. Of note, the activity levels in genotype-matched males vs. females showed differences for both the 1^st^ and 2^nd^ CW introduction (**Sup. Fig. 4A-B**), emphasizing the need for sex-specific analyses. Essentially, activity levels suggest that at 8w post-tmx, mutant mice were just as motivated and physically capable of running as WT mice.

**Figure 2.**
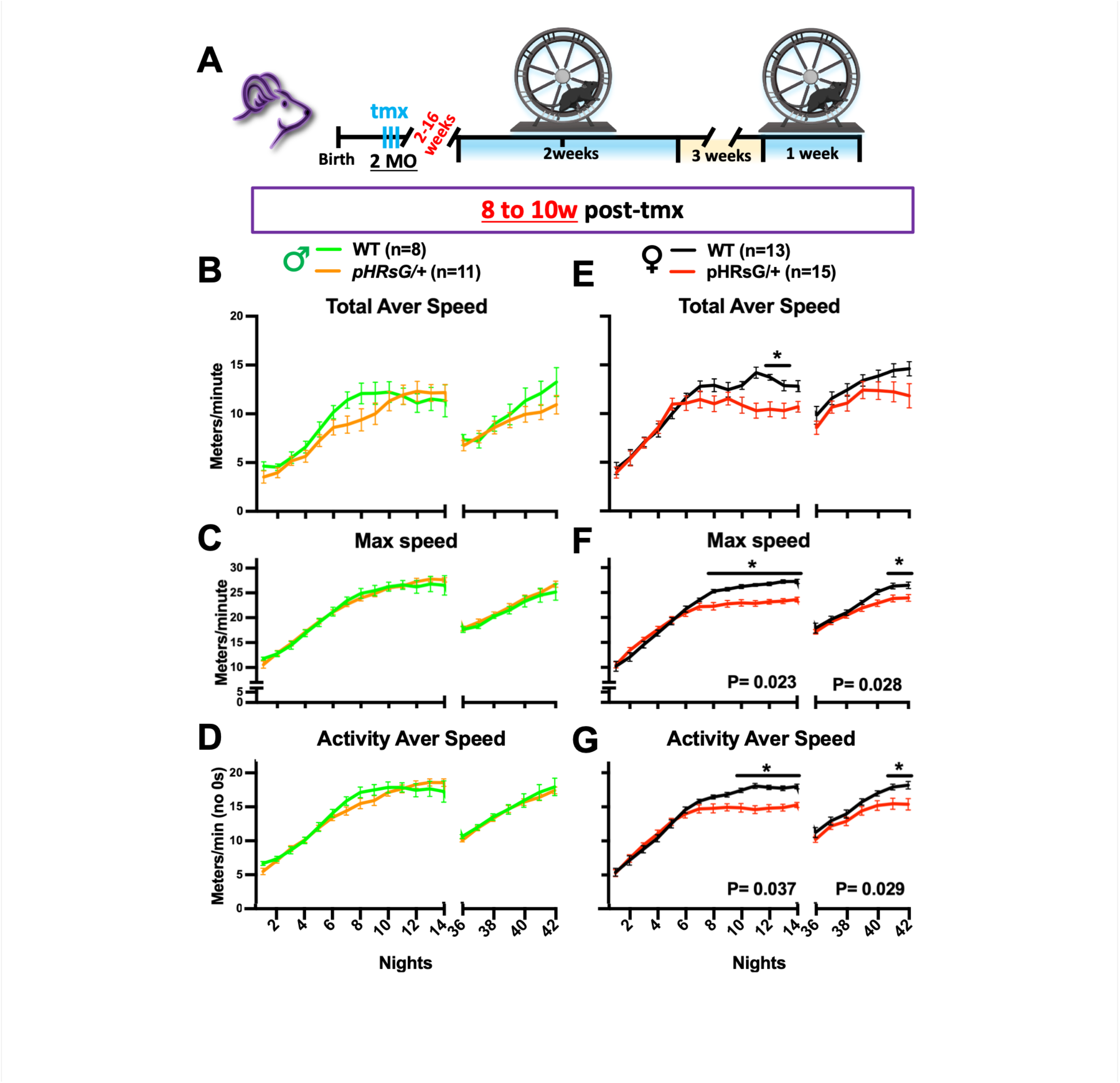
Learning but not Memory Issues in pHRsG/+ Females at 8w Post-Mutation. **A)** General complex wheel (CW) test schedule: 2-month-old (2MO) mice are treated with tamoxifen (tmx; cyan) and 2 to 16 weeks (2-16w) later are introduced to CWs for 2 weeks. After a 3w-break without wheels, mice are reintroduced to CWs for 1 week. **B-I)** The plots for nightly (12h dark cycle) values of CW parameters in WT male (green) and female (black), as well as pHRsG/+ male (orange) and female (red) mice are shown; total average speed (**B, E**; TAS), maximum speed (**C, F**; MaxSpd), and activity average speed (**D, G**; AAS). Statistical comparisons with significant differences show their P values under the plots for the 1^st^ (N1-14) or 2^nd^ (N36-42) introductions (two-way ANOVA test; <u>genotype as source of variation</u>); the curves for MaxSpd (P=0.023 and P=0.028 for 1^st^ and 2^nd^ introduction) and AAS (P=0.037 and P=0.029, respectively) were significantly lower in female pHRsG/+ vs. WTs. The “n” per group is shown in the genotype/sex color code. Bonferroni’s post-hoc test indicates significantly lower values in pHRsG/+ females for TAS in N12-13; MaxSpd in N8-14 and N35-36, and AAS in N9-N14 and N41-42. (**\***p<0.05 for all parameters). Labels in the X and Y axes are shared for panels in the same column or row.

**Figure 3.**
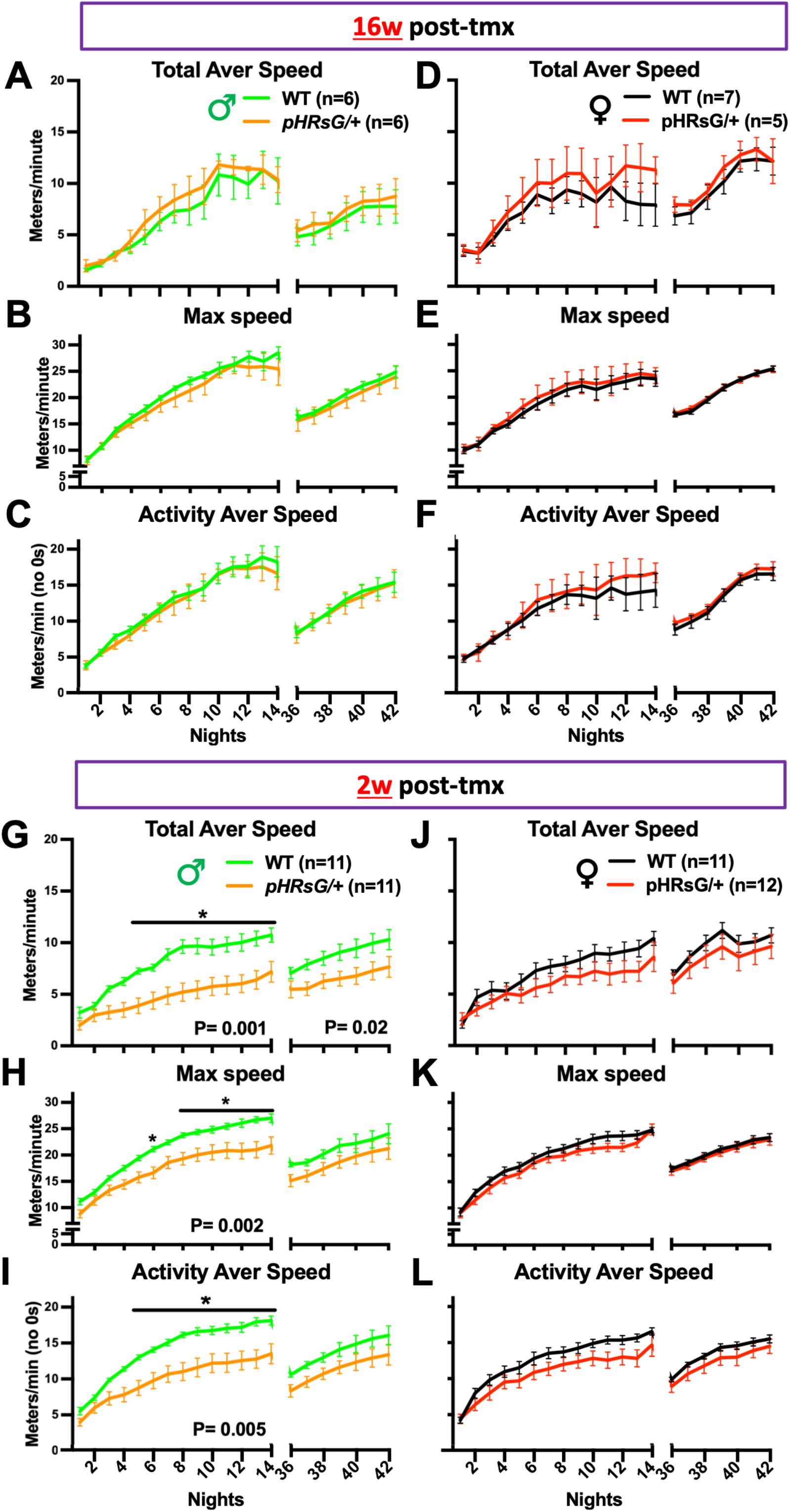
Learning Issues in Male at 2w, but Normal Learning in Male and Females at 16w Post-Mutation. Plots for nightly values of CW parameters in WT female (black) and male (green), as well as pHRsG/+ male (orange) and female (red) mice, tested at 16 weeks (**A-F)** or 2 weeks (**G-L**) post-tmx; total average speed (**A, D, G, J**; TAS), maximum speed (**B, E, H, K**; MaxSpd), and activity average speed (**C, F, I, L**; AAS). WT vs. pHRsG/+ comparisons with statistically significant differences show P values under the plots for the 1^st^ (N1-14) or 2^nd^ (N36-42) introduction to CWs (two-way ANOVA test, with <u>genotype as source of variation</u>, the “n” per group is shown in the genotype/sex color code); no differences between genotypes were found at 16w post-tmx (**A-F**); the curves for TAS (**G**; P=0.001 and P=0.02 for 1^st^ and 2^nd^ CW introduction), MaxSpd (**H**; P=0.02 for 1^st^ introduction) and AAS (**I**; P=0.005 for 1^st^ introduction), were lower in pHRsG/+ vs. WT male mice. Individual nights with significantly lower values in pHRsG/+ mice vs. WTs are also shown (Bonferroni’s post-hoc test **\***P<0.05 for all parameters). Labels in the X and Y axes are shared for panels in the same column or row.

**Figure 4.**
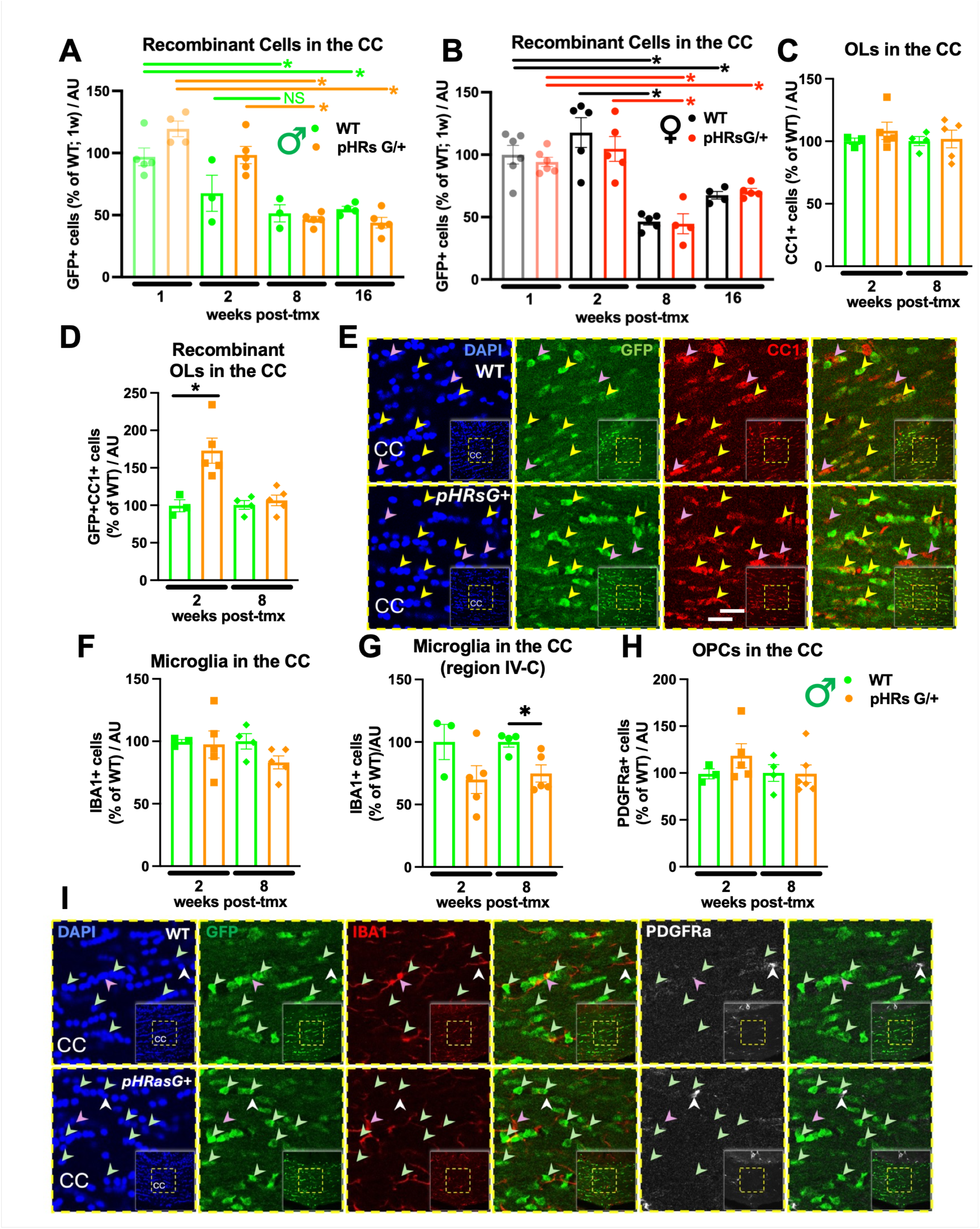
Cell Autonomous and Non-Cell-Autonomous Effects in the CC of pHRsG/+ Mice. **A-B**) The abundance of GFP+ recombinant cells in the corpus callosum (CC) of WT (black) and pHRsG/+ (red) female mice, as well as WT (green) pHRsG/+ (orange) males, at 2w, 8w, and 16w post-tmx, as normalized to EGFP+ cells in WTs at 1w post-tmx (% of WT; data from Fig. 1E; faded bars) is shown. No statistical differences between WT and pHRsG/+ EGFP+ cell abundance were found at any time point. Comparisons for <u>male</u> WTs (green bar/asterisks), and pHRsG/+ (orange bars/asterisks), as well as <u>female</u> WTs (black bars/asterisks) and pHRsG/+ (red bars/asterisks), are shown. *P<0.05 Student’s *t* test; NS: non-significant. **C-E**) The percentage of immunostained CC1+ oligodendrocytes in the CC (**E**; red) is similar in pHRsG/+ and WT mice at 2w and 8w post-tmx (**C**; unpaired Student’s t test, P>0.05); however, the percentage of GFP+CC1+ recombinant OLs at 2w post-tmx is higher in mutants vs. WTs (**D**; P=0.02). Purple arrowheads in **E** show CC1+ cells and yellow arrowheads GFP+CC1+ cells. **F**) The percentage of IBA+ microglia does not show differences in the overall CC (7 regions) of pHRsG/+ mice vs. WTs at 2w (unpaired Student’s t test, P>0.05) or 8w post-tmx; nonetheless, the CC posterior central region (IV-C) shows lower IBA1+ cells in pHRsG/+ mice at 8w post-tmx (**G**, P=0.02). **H)** The abundance of PDGFRa+ OPCs does not show differences in the overall CC of pHRsG/+ mice vs. WTs at 2w or 8w post-tmx as compared with WTs. **I)** Representative image for GFP (green), IBA1 (red), PDGFRa (white) detection and overlays in the CC of WT and pHRsG/+ mice. Dapi was used to stain nuclei and normalize cell densities per arbitrary units (AU). Scale bar = 25μm.

Learning curves for the nightly total average speed (TAS) run on CWs have been utilized to describe new OL-regulated fine motor skills ^26,27,41^. We observed that the TAS curves for male WT and pHRsG/+ mice were not significantly different at 8-10w post-tmx (**Fig. 2B**). In females, although the TAS curve for pHRsG/+ and WT mice overlap for most of the 1^st^ CW introduction, mutants showed lower TAS values during N12-N13 (**Fig. 2E**). Then, we analyzed the nightly maximum speed (MaxSpd) and activity average speed (AAS, average speed of only minutes with activity/night), potentially influencing the TAS changes. Indeed, pHRsG/+ females were not able to reach the MaxSpd achieved by WTs from N8-N14 and subsequently N41-N42, resulting in significantly different curves for the 1^st^ and 2^nd^ CW introductions (**Fig. 2F**). Similarly, pHRsG/+ females had lower curves for AAS, not reaching WT values from N10-14 and N6-N7 (**Fig. 2G**). As expected, pHRsG/+ males did not show significant differences in MaxSpd or AAS as compared with WTs (**Fig. 2C-D**). Consistent with normal motivation and motor capability of mutant mice, all initial CW values for the 1^st^ (N1) and 2^nd^ (N36) introductions did not differ between WTs and pHRsG/+ mice (**Fig. 2B-G, Sup. Fig. 2A-B**). Regarding memory of learned CW skills, the initial values for the 2^nd^ introduction to CWs (N36; after a 3-week break) were significantly higher in both WTs and pHRsG/+ mice as compared with their respective N1 (**Sup. Fig. 3D-I**; N1 vs. N36), but not as high as their N14 (end of 1^st^ CW introduction; N14 vs. N36), indicating that regardless of the genotype, mice retained partial memory of previously acquired skills. Overall, our results indicate that at 8-10 weeks post-tmx, pHRsG/+ females, but not males, exhibit mild deficits in developing fine motor skills, failing to reach WTs levels of MaxSpd and AAS. Whether these phenotypes are progressive is unknown.

### 3.3 Learning Issues are Transient and Sex-Dependent in Mice with OL *HRasG12V* Mutation

Myelin ultrastructural and learning issues develop gradually (over >4 months) in another myelin-focused RASopathy model ^41^; hence, we tested whether CW issues are present in pHRsG/+ males, and/or exacerbated in females, at 16w post-tmx. Activity levels significantly decreased in both male and female, WT and pHRsG/+, mice at 16w as compared with 8w post-tmx (**Sup. Fig. 5A-B**); however, no significant changes in activity levels were detected in pHRsG/+ mutants relative to age/sex-matched WTs at 16w post-tmx (**Sup. Fig. 2C-D**). Importantly, at 16w post-tmx, neither male nor female pHRsG/+ mice showed differences in any CW parameter as compared with WTs (**Fig. 3A-F**), indicating that learning phenotypes observed in pHRsG/+ females at 8-10w post-tmx are transient, while males do not present learning issues at 8w-16w post-mutation. Of note, while males and females showed decreased TAS curves at 16w vs. 8w post-tmx, only females decreased their MaxSpd and AAS (**Sup. Fig. 5F, H**) suggesting sex-independent decrease in motivation or capability to run, but different regulation of acquisition of fine motor skills by sex.

**Figure 5.**
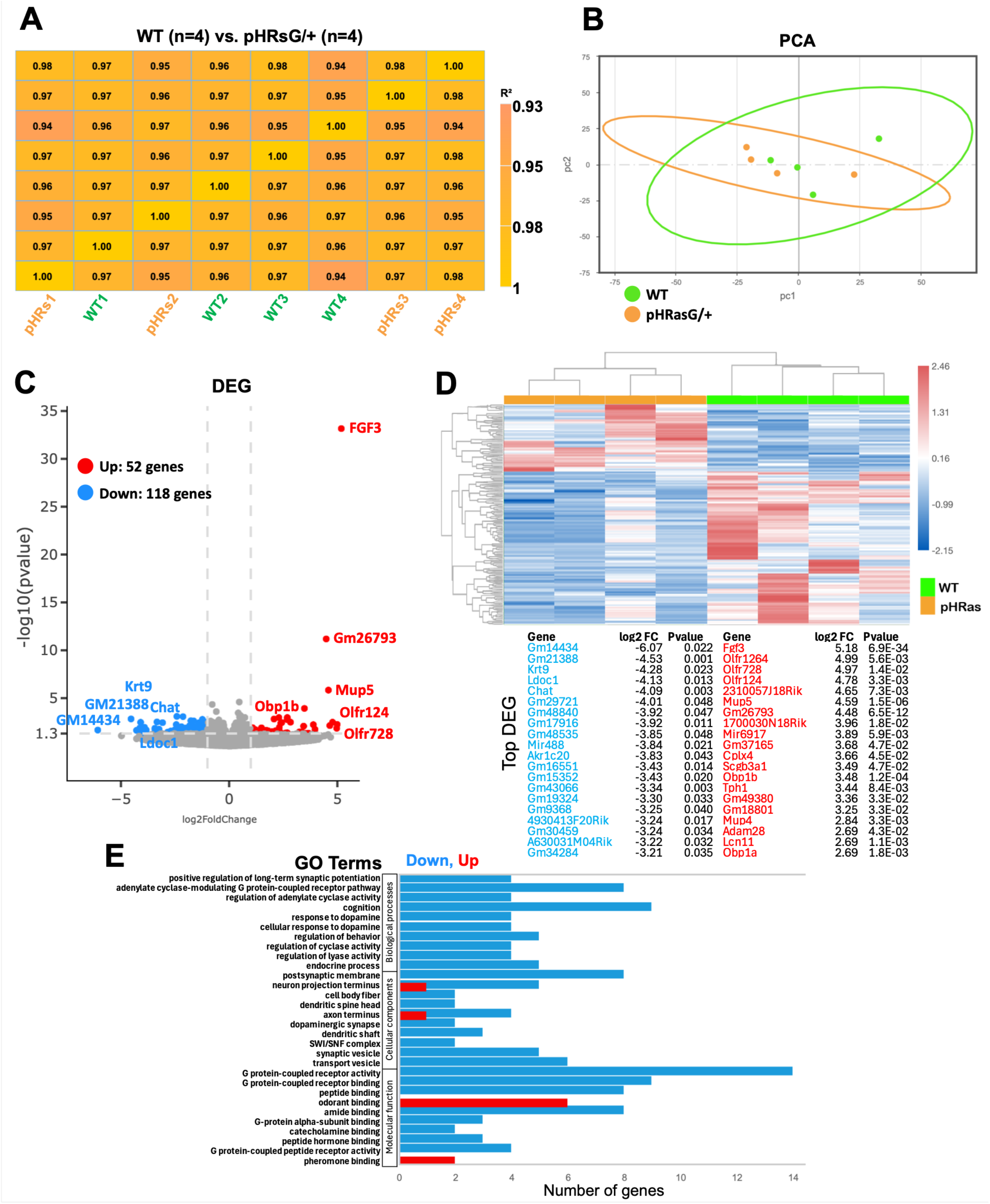
Bulk RNAseq from the ON of pHRasG/+ and WT Male Mice. **A-B**) Pearson’s correlation matrix and principal component analysis (PCA) for the transcriptomes of pHRasG/+ (orange, n=4) and WT (green, n=4) ONs; overall samples show high correlation (r^2^ > 0.94); pHRas samples dispersed between/beyond WTs. **C-D**) Expression level volcano plot and heatmap show up- (red, 52 genes) and down- (blue, 118 genes) regulated genes; differentially expressed genes (DEG) in pHRasG/+ vs. WT ONs (P<0.05; log_2_ fold change >1.0, horizontal line: statistical significance threshold). The top 20 DEG are annotated (**D,** bottom; up regulated in red, downregulated in blue). **E**) Enrichment analysis for biological processes, cellular components, and molecular function GO terms including both up- and down-regulated genes (x axis; number of genes). Down-regulated genes in pHRasG/+ ONs mostly include behavior related GO terms.

The *G12V* mutation of *HRas* drives strong RAS activation and severe CS presentation ^3,48^; hence, we tested whether pHRsG/+ mice develop CW learning phenotypes as early as 2w post-tmx. Interestingly, pHRsG+ males showed robust defects in the CW test as compared with WTs (**Fig. 3G-I**). First, activity levels in pHRsG/+ males were significantly lower during N3-8 but gradually increased until reaching WT values by the end of the 1^st^ CW introduction (**Sup Fig. 2E**). In contrast, the TAS curve in pHRsG/+ males was consistently lower from N5 to the end of the 1^st^ introduction (**Fig. 3G**). The curves for MaxSpd and AAS were also lower in male mutants from N8 and N5 to the end of the 1^st^ introduction, respectively (**Fig. 3H-I**). Consistent with other time points, memory of previously acquired skills seems undisturbed in pHRsG/+ vs. WTs at 2w post-tmx (**Fig. 3G-L**; N36). In summary, pHRsG/+ males develop robust learning issues 2w post-tmx, and females 8w post-tmx; phenotypes seem to be independent of the running activity.

### 3.4 Cell-Autonomous and Non-Cell-Autonomous Responses Upon *HRasG12V* Mutation

Considering the robust and dynamic learning defects in pHRsG/+ males, we analyzed WM cell populations at key time points. To confirm that *HRasG12V* expression in OLs has a functional impact as early as 2w post-tmx, we immunodetected active ERK in the CC; the abundance of total pERK+ and pERK+GFP+ recombinant cells significantly increased in both pHRsG/+ males and females (**Sup Fig. 6**). Next, we evaluated the permanence of GFP+ cells in pHRsG/+ vs. WT mice at 2w, 8w, or 16 weeks post-tmx, as normalized to GFP+ cells in WTs 1w post-tmx (from Fig. 1E). No significant genotype-driven differences were found in males or females (**Fig. 4A-B**); yet pHRsG/+ males showed a trend toward increased GFP+ cells at 1w and 2w post-tmx, causing an apparent slower decrease in recombinant cells, as compared with WTs (**Fig. 4A**; 2w vs. 8w post-tmx). Similar chronological analysis in females showed no differences in GFP+ cells between WT and pHRsG/+ at any time point (**Fig. 4B**). Overall, the transiency and sex-dependent severity of learning phenotypes seem unrelated to the persistence or functionality of mutant cells.

**Figure 6.**
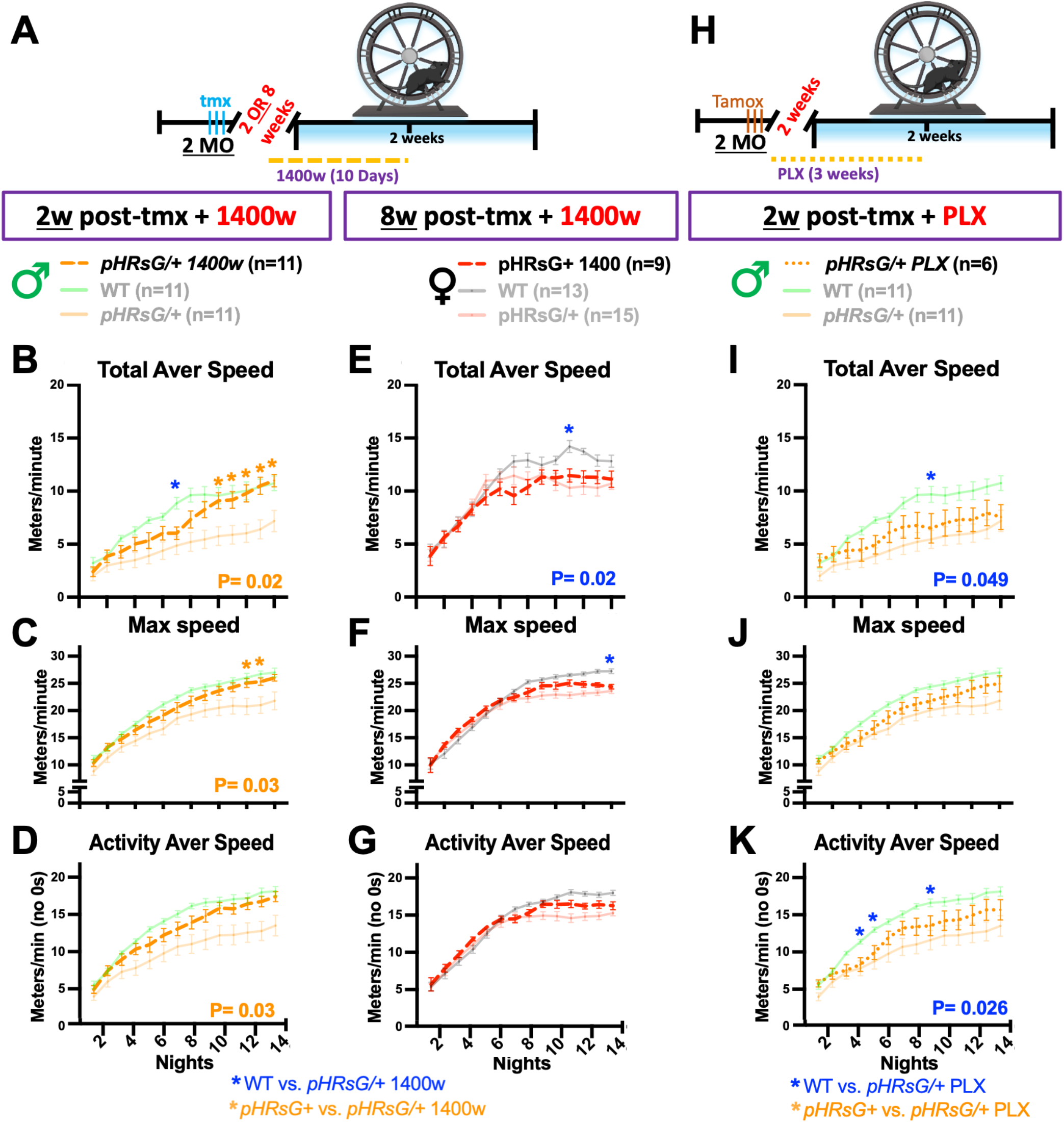
Learning Curves After NOS2 Inhibition and Microglia Depletion in pHRsG/+ mice. **A,H)** treatment schedules: 2-month-old (2MO) mice were treated with tamoxifen (tmx; cyan), and 2w (**B-D, I-K;** males) or 8w (**E-G**; females) later were introduced to CWs for 2 weeks. Mice were treated with 1400w (**A-G**) for 3 days before introduction to CWs and 7 days during the CW test (10 days total; dashed lines). Plots for CW values in 1400w-treated pHRsG/+ male (orange dashed lines) and female (red dashed lines) mice, compared with untreated age/sex matched females (as in Fig. 2, faded solid lines) and males (as in Fig 3G-I, faded solid lines). The P values for significantly different comparisons between 1400w-treated pHRsG/+ mice vs. untreated WT are shown under the plots in blue (two-way ANOVA test, with <u>treatment as source of variation</u>); the TAS for pHRsG/+ females was lower vs. untreated WTs (**E**; P=0.02). Bonferroni’s multiple comparison (blue asterisks) indicates lower TAS for males in N7 (despite no differences in the ANOVA test), and females in N11. Significantly different P values for comparisons between 1400w-treated pHRsG/+ vs. untreated pHRsG/+ males (faded orange line) are shown under the plots (**B-D**; two-way ANOVA test, <u>treatment as source of variation</u>, TAS; P=0.02, MaxSpd; P=0.03, and AAS; P=0.03). Bonferroni’s multiple comparison (orange asterisks) indicates lower TAS (N10-14), and Max speed (N12-13) in 1400w-treated pHRsG/+ males vs. untreated pHRsG/+ mice. **H**) 2MO males were treated with tmx and started on PLX diet for 3 weeks (dotted orange line). **I-K**) Plots for nightly CW values in PLX-treated pHRsG/+ males (orange dotted lines) vs. untreated age-matched males (as in Fig. 3G-I, faded solid lines). Significantly different P values for treated pHRsG/+ vs. untreated WTs are shown in blue (two-way ANOVA test; TAS P=0.049; AAS P=0.026). Bonferroni’s comparison (blue asterisks) indicates lower TAS (N9) and AAS (N4-5, N9) vs. untreated WTs. The “n” per group is shown in the color code. Labels in the X axes are shared for panels in the same column.

Next, to compare time points with different learning phenotypes we quantified OLs at 2w and 8w post-tmx in males and observed that the total amount of CC1+ OLs did not change in pHRsG/+ (**Fig. 4C, E**; percentage of age-matched WTs), yet the density of GFP+CC1+ recombinant OLs was higher in pHRsG/+ mice at 2w, but not 8w post-tmx (**Fig. 4D, E**). This supports the notion of a slower decrease in recombinant cells driven by *HRasG12V*. We then analyzed non-cell-autonomous effects on WM glial cells. The percentage of IBA1+ microglia was not different in the overall CC of pHRsG/+ mice (**Fig. 4-F, I**), yet significant decrease was observed at 8w post-tmx in the posterior-central CC region (**Fig. 4G**, region IV-C). Finally, we did not observe significant changes in PDGFRa+ OPC numbers in the total CC of pHRsG/+ mice (**Fig. 4H**), but a significant decrease was detected in the anterior central region (**Sup. Fig. 1F**, region I-C). In general, cellular mechanisms influencing the development and later extinction of learning issues in pHRsG/+ might include slightly longer persistence of recombinant cells, as well as microglia and OPC responses.

### 3.5 Bulk mRNA From the Optic Nerve

The optic nerve (ON) is rich in OLs and has been used as a readout of transcriptional changes upon OL mutations ^54,55^; hence, we performed bulk ON RNAseq from WT and pHRsG/+ males 2w post-tmx. Unsupervised gene expression clustering (4 ONs/genotype) showed high correlation among samples (Pearson’s > r^2^ 0.94) and principal component analysis indicated modest transcript variance between genotypes (**Fig. 5A-B**), likely because of the short period after *HRas* mutation. Nonetheless, analysis of differentially expressed genes (DEG; WT vs. pHRs; P<0.05; log^2^ fold change >1.0) indicated upregulation of 52 and downregulation of 118 genes (**Fig. 5C-D**). The top DEGs in pHRsG/+ are not typical indicators of OL dys/function and the highest upregulated gene, Fgf3, is not associated with specific roles in OLs, yet the FGF family regulates OL function ^56^ and OPC responses to compromised myelin ^57^. Other top upregulated genes are associated with autoimmune disease (**Obp1a** ^58^), cytokine biology (***Scgb3a1*** ^59^), immune and inflammatory pathways (**Cplx4** ^60^) including in microglia (**Chat** ^61^, **ADAM28** ^62^) and immune OLs (**ADAM28** ^63^). Interestingly, functional GO enrichment analysis indicated that downregulated genes/terms are associated with behavior control including long-term synaptic potentiation, cognition, response to dopamine, regulation of behavior, regulation of synaptic plasticity, learning or memory (biological processes); postsynaptic membrane, dendritic spine and shaft, axon terminus, synaptic vesicle (cellular components), and G protein-coupled receptor activity/binding, odorant/pheromone binding (molecular functions) among others (**Fig. 5E)**. Overall, *HRasG12V* mutation induces in the short-term responses associated with immune response and networks with negative impact on high order brain function.

### 3.6 NOS2 Inhibition Rescues Learning Issues in pHRsG/+ Males

Short-term treatment with broad NO synthase (NOS) isoform inhibitors rescues myelin structural phenotypes in pHRsG/+ and another CS model ^49,64^. Considering the putative microglial activation phenotype (**Figs. 1F-G, 4F-G**) and inflammation-immune transcriptomic signatures (**Fig. 5 C-E**), we tested whether the NOS2-specific inhibitor 1400w (Kd=7 nM; with OL protective roles ^65,66^) reverts learning phenotypes in pHRsG/+ mice. Mice were IP injected with a safe dose of 1400w [2mg/kg ^67^] one-week post-tmx for 14 days; however, both WT and mutant mice became lethargic, and necropsy showed liver damage (**Sup. Fig. 7**); the genetic background of our strain could contribute to such susceptibility. To minimize negative/acute effects of the drug, pHRsG/+ mice were subcutaneously treated with a lower dose of 1400w (0.2mg/kg per day), starting three days before and seven days during the 1^st^ introduction to CWs (**Fig. 6A**, 2w post-tmx in males and 8w post-tmx in females); no negative effects of the drug were observed in the liver or other organs. Remarkably, 1400w treatment rescued abnormal CW parameters in pHRsG/+ males. In detail, although initial activity levels in 1400w-treated pHRsG/+ mice were similar as in untreated mutants, values progressively increased and matched those in untreated WTs, resulting in no significant differences between CW curves of 1400w-treated pHRsG/+ vs. untreated WTs (**Sup. Fig. 2G**), except for lower activity in N7 (Bonferroni’s post hoc test). Moreover, significantly different activity was observed when comparing 1400w-treated mutants vs. untreated ones (**Sup. Fig. 2G)**. Similarly, the curves for TAS were not significantly different between 1400w-treated pHRsG/+ and untreated WT mice (**Fig. 6B**), yet values for N7 were lower, suggesting a link with the above N7 activity levels. Remarkably, MaxSpd and AAS curves for 1400w-treated pHRsG/+ males overlapped with those of untreated WT mice (**Fig. 6C-D**). To better describe the extent of learning rescue, CW values from 1400w-treated pHRsG/+ mice were compared with untreated pHRsG/+ mice. The curves for TAS, MaxSpd, and AAS were significantly higher in treated pHRsG/+ males as compared with untreated mice, with robust differences starting at N10 for TAS, and N12-13 for MaxSpd and AAS (**Fig. 6B-D**). Of note, NOS2 inhibition caused detrimental effects on male WT mice, as the activity level, TAS, MaxSpd, and AAS values were significantly lower in 1400w-treated WT mice as compared with untreated WTs at 2w post-tmx (**Sup. Fig. 8A-D**). On the other hand, 1400w-treated WT females showed lower activity and TAS, but not MaxSpd or AAS vs. WTs, at 8w post-tmx (**Sup. Fig. 8E-H**). As previously suggested ^41^, these results indicate that any imbalance in NO signaling (increase or decrease) can negatively impact brain function.

**Figure 7.**
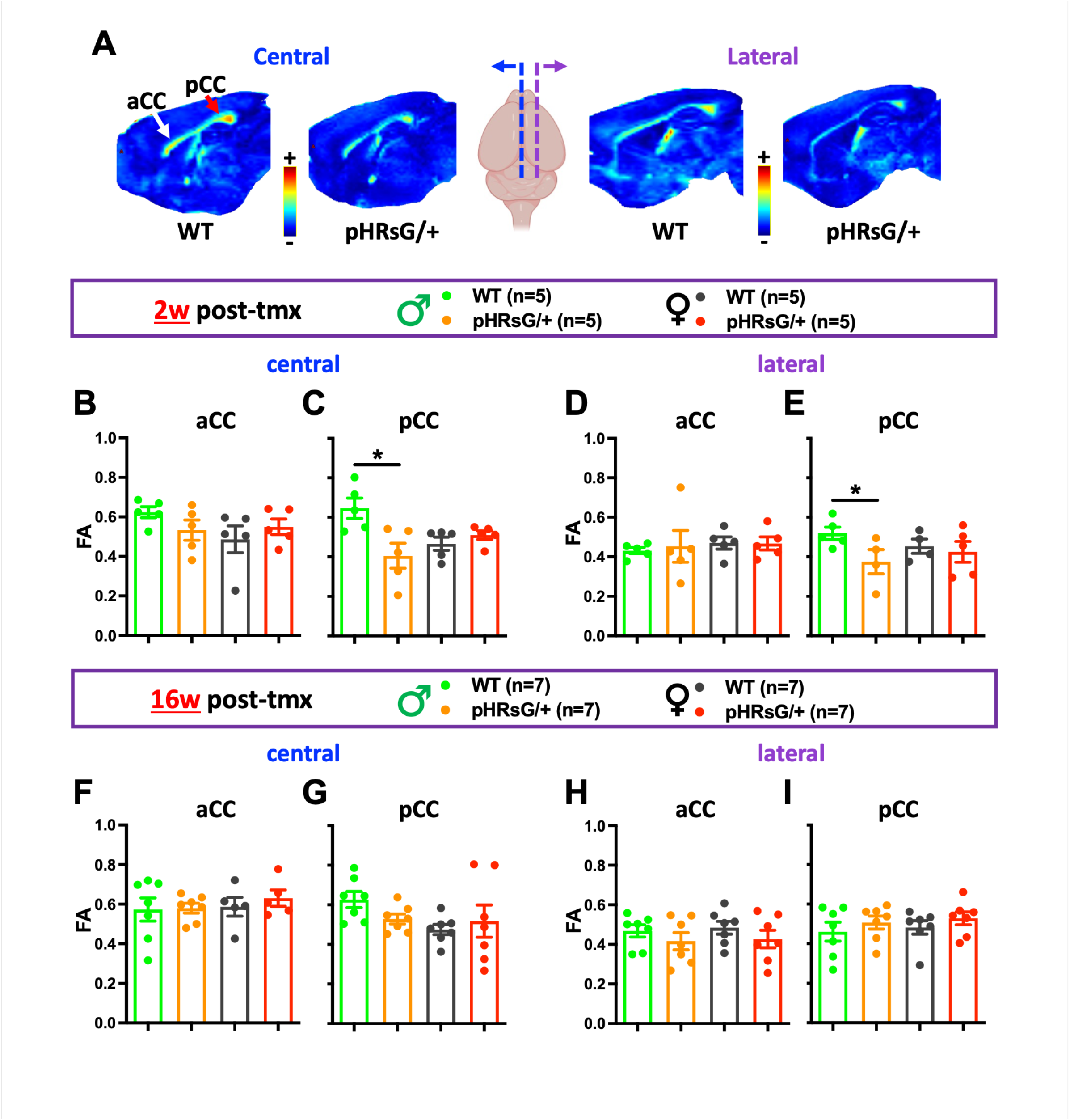
Microstructural Abnormalities in the Corpus Callosum of Male pHRasG/+ Mice at 2w, but not 16w, After *HRas* Mutation. **A**) Representative sagittal plane pictures of diffusion imaging analyses (DTI-MRI), showing fractional anisotropy (FA) heat maps in the anterior (aCC, white arrow) and posterior (pCC, red arrow) corpus callosum regions. The brain cartoon indicates the central-lateral CC regions analyzed (central: blue-dashed line; lateral: purple-dashed line). **B-E)** FA values 2w post-tmx were graphed for male (green) and female (black) WT mice, as well as male (orange) and female (red) pHRsG/+ mice, in the four regions depicted in A. Significantly lower FA values were observed both in the central and lateral pCC of pHRsG/+ males as compared with WTs (unpaired Student’s t-test; P=0.009 and P=0.031, respectively). The “n” per group is shown in the genotype/sex color code. **F-I)** FA values at 16w post-tmx are shown for the same CC regions assessed in B-E. No significantly different FA values were detected for any CC region in pHRsG/+ vs. WT, males or female mice.

The effect of 1400w was also tested in pHRsG/+ females at the peak of their learning issues (8w post-tmx); significantly lower activity curves and lower values in N1 and N7 for pHRsG/+ females as compared with untreated WTs (**Sup. Fig. 2H**) were found. This contrasts with the activity rescue elicited by 1400w in Ms; however, females did not show lower activity at any time point (**Sup. Fig. 2B**). The TAS curve was still significantly lower in pHRsG/+ females vs. untreated WTs (**Fig. 6E**), suggesting that 1400w does not rescue the learning phenotype; however, the MaxSpd and AAS curves were not different (**Fig. 6 F-G**). The CW curves for 1400w-treated pHRsG/+ females were not different vs. untreated pHRsG/+ females (**Fig. 6 E-G**). These inconclusive results could relate to the impact of 1400w on activity levels and TAS also found in WT females (**Sup. Fig. 8, E-F**), without impacting MaxSpd or AAS (**Sup Fig. 8, G-H**). Overall, NOS2 control had a clear positive impact on learning phenotypes in pHRsG/+ males, yielded inconclusive results in mutant females, and negatively impacted both male and female WT mice.

NOS2 increases in *HRasG12V* mutant OLs one-month post-tmx ^64^ but transcripts did not increase at 2w post-tmx in pHRsG/+ mice (**Fig. 5C-D**). To narrow potential sources of increased NO influencing learning phenotypes, we used the agent PLX3397 (PLX) to deplete microglia. Male pHRsG/+ and WT mice were fed ad libitum with a diet containing a safe/effective amount of PLX [290 mg/kg, ^68^], for three weeks starting the day of tmx treatment (**Fig. 6H**); treatment included 7 days of the 1^st^ introduction to CWs. A slight reduction (4.1%) in mouse body weight was detected at the start of the CW test, but no obvious negative effects were observed. Activity levels in PLX-treated pHRsG/+ mice were similar to those in WTs, only showing a trend toward decrease by the end of the 1^st^ CW introduction (**Sup. Fig 2I**). The curves for TAS and AAS were still significantly lower in PLX-treated pHRsG/+ mice as compared with untreated WTs (**Fig. 6I, K**, including lower values in N7 for TAS, and N4-5, N10 for AAS) indicating that PLX does not rescue these parameters. In contrast, the MaxSpd curve did not show differences between treated pHRsG/+ and untreated WT mice (**Fig. 6J**). On the other hand, no impact of PLX treatment was observed for any CW parameter when comparing PLX-treated vs. untreated pHRsG/+ mice (**Fig. 6I-K**). Finally, PLX treatment in WT mice did not cause changes in activity, TAS, or AAS, but caused a modest decrease in MaxSpd, as compared with untreated WT (**Sup. Fig. 8I-L**). Overall, our results suggest that microglial NOS2 is not the only or main source of the NO mediating learning issues in pHRsG/+ males; whether microglia function is necessary for the compensatory mechanisms observed later, remains to be studied.

### 3.6 dMRI Analysis Reveals Abnormal WM Microstructure Concomitant with Transient Learning Issues in Male pHRsG/+ Mice

Next, we tested whether OL-related structural defects, can be detected at the time of learning deficits in pHRsG/+ mice. To explore a clinically relevant marker of myelin microstructure changes (provided that myelin decompaction assessment in patients is impractical), commonly available diffusion MRI (dMRI) methods were used. We collected and fixed brains of male and female WT and pHRsG/+ mice at 2w post-tmx and subjected them to dMRI analyses; four central-lateral and anterior-posterior regions were analyzed (**Fig. 7A**). Remarkably, quantification of fractional anisotropy (FA) showed significant decreases in both the central and lateral regions of the posterior CC (pCC) of pHRsG/+ males, but not in the anterior CC (aCC), as compared with WT mice (**Fig. 7B-E**). These results suggest regional heterogeneity in the impact of *HRas* mutation on myelin structure 2w post-tmx, despite no regional differences in recombination (**Figs. 1E, 4A**). In correlation with the absence of learning phenotypes in pHRsG/+ females at 2w post-tmx, their FA values did not show significant differences in any CC region analyzed, as compared with WT females (**Fig. 7B-E**). These results substantiate sex-dependent differences for the impact of *HRas* mutation in OLs. If decreased FA values are related to learning issues, we would expect normalization of FA values at a time point when learning is back to normal. Indeed, we observed that at 16w post-tmx the FA values of pHRsG/+ males, and females, were not significantly different in any of the regions inspected, as compared with WTs (**Fig. 7F-I**). Overall, shortly after inducing *HRasG12V* mutation in OLs of male mice, the WM microstructure of the posterior CC shows reduced integrity that resolves by 16w post-tmx.

## 4. DISCUSSION

The impact of *HRas* mutation on adult OL biology and its links with neurological issues remain poorly explored. Replacing the *HRas* gene with the gain of function *HRasG12V* mutant version in adult myelinating cells (pHRsG/+ mice), hyperactivates Erk-NO-Notch signaling and causes ultrastructural issues ^49,64^; yet no brain function consequences are known. Here, we used pHRsG/+ mice to assess learning and associated cell-autonomous/non-cell-autonomous, WM microstructural, transcriptional, and NO-mediated mechanistic phenotypes, at various time points after mutation. We detected sex-dependent abnormal learning in the CW test; males showed robust issues at 2w post-tmx while females showed mild issues at 8w, and learning normalized by the week 8 and 16 after mutation, respectively. Compensatory mechanisms involving the OL lineage and microglia likely contribute to the extinction of phenotypes. A key molecular mediator of phenotypes is NOS2 activity, as its inhibition rescued learning; hence, NOS2 emerges as a potential therapeutic target in RASopathy neuropathology. In synchrony with the peak and loss of learning phenotypes, male mutants showed abnormal WM microstructure (decreased FA) at 2w but not at 16w post-tmx. Our OL-inducible *HRas* mutation model suggests that abnormal signals might cause chronic issues in germline CS models or in patients, in which *HRas* mutation is ubiquitous and constitutive/developmental. Overall, the view that adult brain dysfunction in genetic diseases is permanent has been previously challenged in RASopathies ^55,69–77^; our study supports this idea and suggests that learning phenotypes linked to abnormal OL biology can be modified.

### 4.1. The pHRsG/+ Model; Discoveries and Limitations

Various CS models mimic patient features including facial dysmorphia; nonetheless, whether models mimic neurological phenotypes is less clear ^69,78,79^. For better CS modeling, we changed from a transgenic *HRasG12V* system ^64^ to a “flox and replace” (FR) model that switches *HRas* for FR-*HRasG12V* in its endogenous locus ^49,50^. Mice with germline FR-*HRasG12V* mutation show MAPK hyperactivation, tumor predisposition ^50,80^, and abnormalities in the corpus callosum ^81^, which mimics the CS patient phenotype. To study mature OL contribution to neurological issues, we induced heterozygous *HRasG12V* mutation in Plp+ cells of adult mice. As in previous studies using the PlpCreER driver in adults ^41,55^, we observed efficient, specific, and sex-unbiased recombination in OLs, yet quick responses in non-targeted OPCs, astrocytes, and microglia were found. Interestingly, demyelination models displaying CW learning issues ^43^ also show microglia and NO responses ^82,83^, resembling our observations in pHRsG/+ mice; therefore, it is probable that abnormal myelin (not absent) triggers compensatory mechanisms similar to those during demyelination. It is also probable that CW-stimulated OPC differentiation ^26^ is less efficient during compensatory responses in pHRsG/+ mice in the short-term, but not in the long-term. While the compensatory machinery restoring brain function in pHRsG/+ is intriguing, we prioritized exploring the short-term mechanisms involved in robust learning defects, mainly in males. As our model targets primarily mature OL, mechanisms are not intrinsic to known defective OPC differentiation precluding CW learning ^26,27^. Instead, we propose that hyperactive RAS/MAPK signaling in OLs increases diffusion of NO, which signals to other cell types including microglia, compromises myelin integrity or other functions of mutant and neighbor OLs, finally impairing CW learning (**Sup. Fig. 9**). The NO-mediated abnormal myelin g-ratio and decompaction reported in pHRsG/+ mice ^49^ are clear candidates for defective learning cellular mechanism; however, these myelin defects are sustained for at least 6 months post-mutation and found in both male and female mutants ^49,64^, which conflicts with the transient and sex-dependent features of learning deficits in pHRsG/+ mice. Therefore, although myelin decompaction was associated with reduced FA in patients and models of the RASopathy NF1 and Noonan Syndrome ^84,85^, it cannot solely explain the behavior phenotypes of pHRsG/+ mice. So, we hypothesized that myelin decompaction and other OL defects could be detected by broad WM analyses like dMRI; the decreased FA observed at 2w post-tmx in the CC of pHRsG/+ males, but not of females, supports this notion, and the normalization of FA 16w post-tmx further supports the idea. What other OL functions could contribute to learning phenotypes? It was reported that increased inflammation negatively correlates with FA values in the CC of patients with depression ^86^, and our histological and transcriptomic analyses in pHRsG/+ mice point toward inflammation/immune responses, known to regulate OL biology ^33,34^. Other OL roles potentially compromised, include OL metabolic functions supporting axons ^31,32^. Overall, further detailed cellular-molecular analyses associated with defective microstructure is needed identifying specific mechanisms for compromised learning.

Our study has the limitation of targeting *HRasG12V* mutation only in adult oligodendrocytes, in a *HRas* WT context. Future studies must analyze mutations in broader cell populations (including the germline) and time points; nonetheless, the present study unveils important principles of OL *HRas* mutation; 1) abnormal signaling from pre-existing OLs leads to defective learning, 2) NOS2-driven compromised learning suggests a mechanism with therapeutic potential, 3) induced abnormalities in mature OLs/myelin trigger compensatory responses to restore brain function, 4) responses to a CS-causing mutation can be sex-depended, and 5) there is a correlation between compromised WM microstructure and defective learning in our CS model. Overall, our study supports key roles of abnormal mature OLs and/or myelin in the neuropathophysiology of CS.

### 4.2. CW Learning is Compromised by *HRas* Mutation in Mature OLs

Mice carrying transgenic *HRasG12V* in differentiating OLs showed increased locomotion and hypersensitivity to startle behavior but did not show learning/memory issues in the Morris water maze, a test greatly linked to hippocampal neuronal function ^64^; hippocampal-dependent tasks may not capture OL-regulated learning deficits. Instead, fine motor learning is regulated by brain machineries involving the WM; responses of the OL lineage in the CC are necessary for proper CW learning ^26,27^. In our study, the CW test was used, particularly with focus on the MaxSpd and AAS as main readouts for acquisition of fine motor skills. These parameters are mostly independent of the activity levels, which heavily influence the TAS. Although activity levels can inform motivational issues, also relevant in RASopathies, general activity can be impacted by further factors including energy metabolism, muscle strength, pain, etc. Indeed, pHRas mice showed decreases in MaxSpd and AAS in the short-term (2w post-tmx), independently of decreased activity, thus supporting central roles of brain dysfunction, rather than the involvement of other body systems. Moreover, the involvement of compromised WM microstructure in the observed learning phenotypes is supported by decreased FA in males, but not females, at 2w post-tmx and by its normalization at 16w post-tmx.

Interestingly, despite high OL enrichment in the ON, our bulk RNAseq analysis indicated downregulation of classic neuronal function GO terms, mostly relater to behavior. While neuronal RNA transits by ON axons ^87,88^, its relative low abundance raises the question of glial participation in such processes. This possibility is emphasized, as typical or RASopathy-specific ^55^ genes for compaction showed no expression level changes. Overall, the mechanism for impaired mature OLs to directly (via regulation of axonal function) or indirectly (via third party responses) impact learning, merit further research.

### 4.3 NO as Key Mediator of Phenotypes in pHRasG/+ Mice

NOS2 was upregulated in the ON of a CS transgenic model ^64^ and pHRsG/+ mice showed increased number of OLs, but not OPCs, with NO signals one-month post-tmx, as measured by flow cytometry ^49^. We did not observe NOS2 upregulation in our RNAseq analysis, yet we cannot discard upregulation at the translational or post-translational levels ^89^. Hence, we propose that NO increases via NOS2 around 2w post-tmx and, per its diffusible nature, reaches the extracellular environment and signals to non-mutant neighbor cells, thus potentiating/accelerating its effects (**Sup. Fig. 9**); however, we do not discard that microglia or astrocytes contribute to increased NO, as previously seen after myelin damage. Nonetheless, it was shown that NOS2 increases in cultured OLs following cytokine ^90,91^ or LPS ^92^ exposure. Although NO itself shows low toxicity, at high concentrations it can form strong oxidants lethal for mature OLs ^93–95^. Yet we did not find OL numbers to decrease in pHRsG/+ more than in WT mice; a decrease in recombinant OLs occurred in both genotypes from 1-2w to 8w post-tmx. Nevertheless, future studies should aim to define the threshold number of mutant OLs and NO levels that disrupt OL homeostatic roles in learning. This will be also relevant for drug selection for phenotype rescue experiments. In this regard, we selected the compound 1400w because, i) it has a protective effect on compromised OLs after increased ERK activation and peroxinitrate formation^65^, ii) expression of NOS2 has been found in the OL lineage ^90,92,95^, and iii) 1400w crosses the blood– brain barrier and has been safely tested in humans and lab animal ^96,97^. It is important to highlight that NO signaling is complex and any imbalance (increase or decrease) mediate opposing effects on memory depending on the context ^64,98,99^, as we substantiated with the results of 1400w treatment in WT mice. Finally, the detrimental effects that NO-controlling drugs can cause, such as the liver damage observed with 1400w, must be considered under specific contexts and doses.

### 4.4. Sex-Dependent Phenotypes in pHRasG/+ Mice

Sex was determinant for the rapid development of learning phenotypes in pHRsG/+ mice; however, previously reported phenotypes were not modified by the sex ^49,64^ and myelin decompaction was normalized by various genetic/pharmacological tools with additive effects and in a gene mutation-dose effect. Therefore, we speculate that an interplay among abnormal signaling (modulated by number of mutant OLs, the type of mutation, etc.), structural defects, and compensatory responses including those sex-dependent, shape the learning phenotype. As the myelin proteome is essentially the same between sexes ^100^, we suspect that sex hormone-driven responses to *HRas* mutation are key influencers for sex-dependent learning phenotypes. Interestingly, in a longitudinal study of CS patients, females showed higher brain function than males in domains including communication, daily living skills, and socialization, but no differences in visual cognition or visuomotor functioning were found ^18^. Similarly, caregivers reported maladaptive behavioral concerns specifically for male patients ^18^ and the quality of life scores were higher for females than for males ^101^. Whether sex differences in CS are mimicked in pHRsG/+ mice is a remarkable possibility; meanwhile, our study represents the first experimental evidence for sex-driven phenotypes in CS. Overall, our study suggests that chronically abnormal NO signaling initiated in adult OLs contributes to neurological issues preferentially in males.

## Acknowledgments

We thank Dr. Nancy Ratner (Cincinnati Children’s Hospital; Cincinnati) and Dr. Itamar Kahn (Zuckerman Institute, Columbia, NY) for early conceptual contributions. We thank Juan Ortiz-Retana for assistance during MRI acquisition and Edith Garay for technical support. R-R, M is a bachelor’s student from ENES-UNAM-Juriquilla. We thank the UTRGV-CoHP for support from the SMALL grant program.

## Author Contributions

Conceptualization: López-Juárez A.; Experimental strategy and analysis: López-Juárez A., Cisneros-Mejorado A., Nishiyama A. Winkler A. M.; Investigation: Hernandez D., Martinez C., Figueroa A., López-Lorenzo K., Lopez S., Rodriguez J. Radhakrishnan V., Figueroa A., Cisneros-Mejorado A.; Methodology: Martinez C., López-Lorenzo K., Hernandez D., Figueroa A., Lopez S., Radhakrishnan V., Figueroa A., Rodríguez-Rodríguez M., Cisneros-Mejorado A.; Supervision: López-Juárez A., Nishiyama A.; Writing original draft: López-Juárez A.; Funding acquisition: López-Juárez A.

## Data availability

Additional information required to reanalyze the data reported in this paper is available from the corresponding author upon request.

