## Supplementary figures for "Targeting Nitric Oxide Synthase 2 Reverses Learning Deficits in an Oligodendrocyte-Focused Model of Costello Syndrome"

**This PDF file includes:**

Supplementary Figures 1 to 9

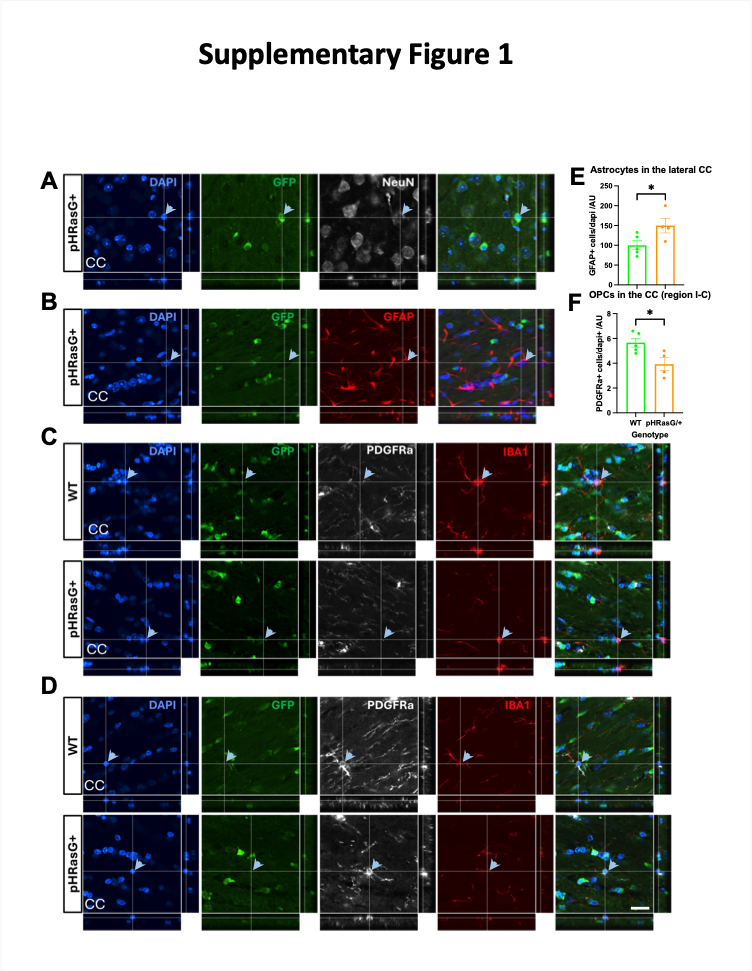

Fig. S1. Recombinant Cells Lack Signals for Astrocyte, OPC, or Neuronal Markers.

Orthogonal views of confocal Z stack images from cortical (A) or corpus callosum (CC; B-D) mouse brain sections immunostained for GFP+ recombinant cells (A-D), NeuN+ neurons (A), GFAP+ astrocytes (B), PDGFRa+ OL progenitor cells (OPCs), and IBA1+ microglia (C-D). Dapi staining was used to identify nuclei. A) The arrow shows a GFP+ cell body in close apposition with a NeuN+ neuron, and the orthogonal views indicate absence of overlap between GFP and NeuN signals (white panel); the GFP+ cell likely corresponds to a recombinant satellite OL. B) example of an GFAP+ astrocyte (arrow) lacking signals for GFP. C) Orthogonal views of the CC of WT and pHRsG/+ brains showing microglia cell bodies (IBA1+; red panel; arrow) lacking signals for GFP. Note the condensed microglia cell body in the brain of the pHRsG/+ mouse. D) Orthogonal views of the CC of WT and pHRsG/+ brains showing OPC cell bodies (PDGFRa+; white panel; arrow) lacking signals for GFP. Quantification of GFAP+ cells (E) shows significantly higher percentage of astrocytes in lateral CC regions (I-B to III-B; in Fig. 1C) of male pHRsG/+ mice, as compared with WTs (unpaired Student’s t test, P=0.048; WTs n=5 and pHRsG/+ n=4). Quantification of PDGFRa+ cells (F) indicates a significantly lower number of OPCs in the anterior central CC (region I-C) of pHRsG/+ mice (unpaired Student’s t test, P=0.025; WTs n=5 and pHRsG/+ n=4). Scale bar = 25 μm.

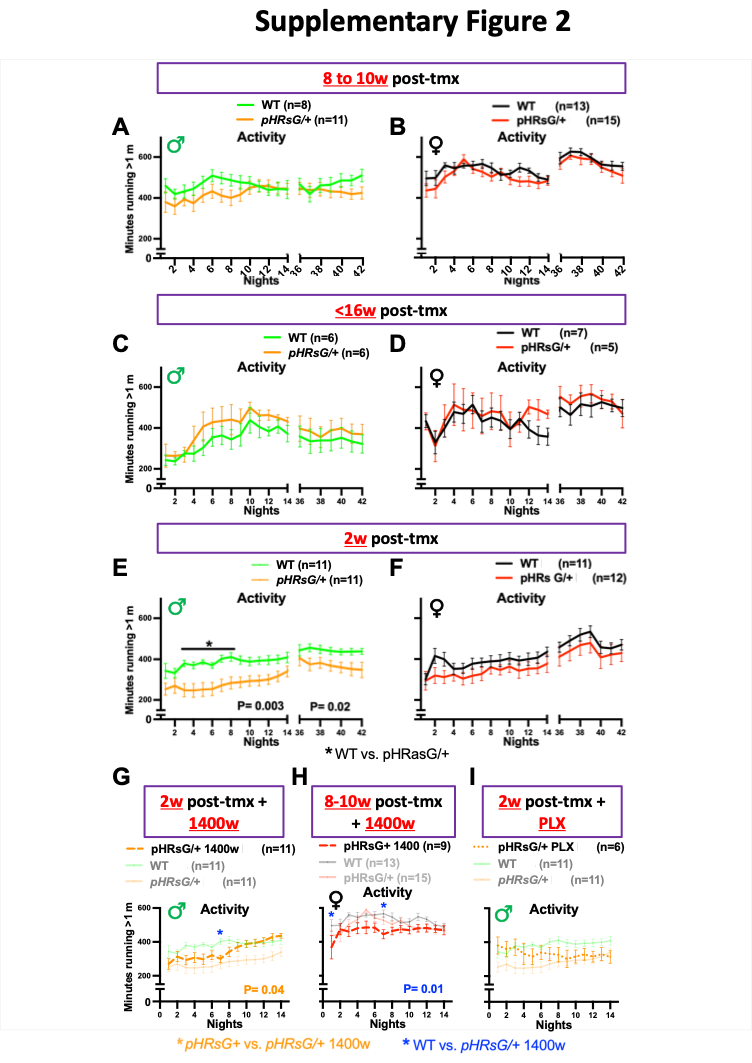

**Fig. S2. CW Activity Levels at Different Time Points Post-Mutation and After Pharmacological Treatments.**

A-I) Plots for CW activity levels (minutes running >1 meter/12h dark cycle) for WT female (black) and male (green), as well as for pHRsG/+ male (orange) and female (red) mice, at the indicated time points post-treatment (purple boxes). The P value for statistically significant differences between WT and pHRsG/+ mice are shown under the plots of the 1^st^ (N1-14) and 2^nd^ (N36-42) CW introductions (two-way ANOVA test with genotype as source of variation; Bonferroni’s post hoc test; *P<0.05 for individual night [N] comparisons). Activity levels are lower in pHRsG/+ males vs. WTs at 2w post-tmx (E, P=0.003 and P=0.02 for the 1^st^ and 2^nd^ introduction), including lower values from N3-8 (*black). G-H) Activity levels for 1400w-treated pHRsG/+ males (2w post-tmx; orange dashed line) and females (8-10w post-tmx; red dashed line) are compared with age/sex-matched untreated mice (same as E and F, respectively; faded solid lines). There are different activity levels in 1400w-treated pHRsG/+ (n=11) vs. untreated pHRsG/+ (n=11) males (G; two-way ANOVA test, treatment as source of variation; P=0.04). No curve differences were found in untreated WT vs. 1400w-treated pHRsG/+ males; however, the there is a difference in N7 (blue *P<0.05). Activity was different between 1400w-treated pHRsG/+ females (n=13) vs. untreated WTs (n=9; two-way ANOVA, P=0.01) and there were differences in N1 and N7 (blue *P<0.05). I) Activity in pHRsG/+ males treated with PLX3397 (PLX; 2w post-tmx; orange dotted line) are compared with untreated males (same as E; faded solid lines). Labels of Y axes are shared in A-B, C-D, E-F, and G-I panels.

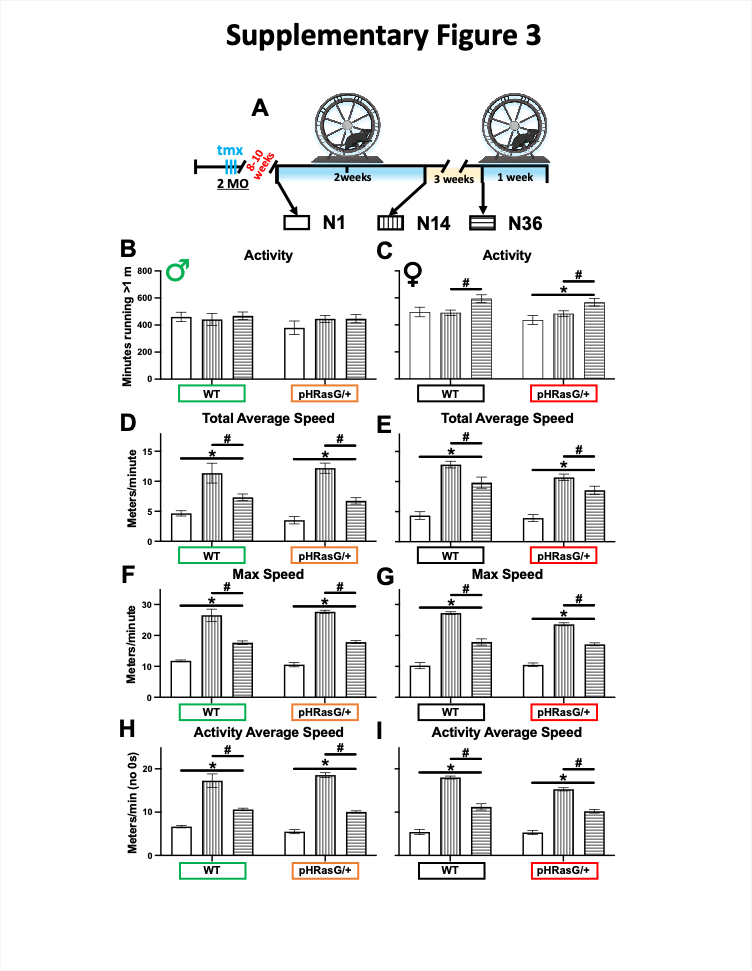

**Fig. S3. *HRasG12V* Mutation in OLs does not Disrupt Memory of Fine Motor Skills.**

A) Experimental CW schedule showing tmx treatment, waiting period (8-10w, red), night 1 (N1, empty box), N14 (end of 1^st^ CW introduction; box with perpendicular lines), and N36 (start of 2^nd^ CW introduction; box with horizontal lines), which are compared in B-I. Graphs for CW activity levels (B-C), total average speed (D-E), max speed (F-G), and activity average speed (AAS; H-I), in male (B, D, F, H) and female (C, E, G, I) mice. The genotypes for WT males (green) and females (black), as well as pHRsG/+ males (orange) and females (red) are shown in the X axis. Unpaired Student’s t test was used to compare CW parameters in N36 vs. N1 (start of vs. end of the 1^st^ CW introduction; *P<0.05) and N36 vs. N14 (end of 1^st^ CW introduction vs. start of the 2^nd^; ^#^P<0.05).

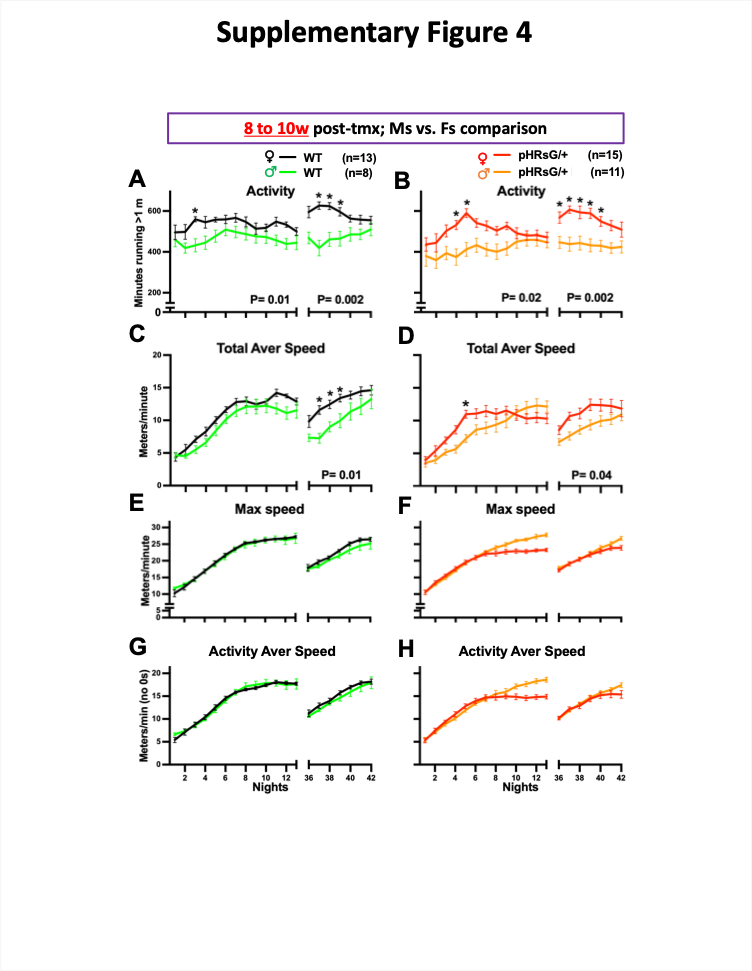

**Fig. S4. Sex is a Determinant of CW Activity Levels and Average Speed.**

Plots for nightly values of CW parameters in WT female (black) and male (green), as well as in pHRsG/+ male (orange) and female (red) mice. Male vs. female comparisons for CW activity levels (A-B), total average speed (C-D; TAS), maximum speed (E-F; MaxSpd), and activity average speed (G-H; AAS) are shown. Statistically significant P values are shown under the plots (two-way ANOVA test, with sex as source of variation); specifically, activity levels in male WT (A; P=0.01 and 0.002 for 1^st^ and 2^nd^ CW introduction) and pHRsG/+ (B; P=0.02 and P=0.002) mice are lower as compared with females; TAS is lower during the 2^nd^ CW introduction in WT males (P=0.01) and mutant males (P=0.04). Bonferroni’s post-hoc test (*P<0.05 for all parameters) indicates significantly lower activity levels in WT males vs. females (N2, N37-39) and pHRsG/+ males vs. females (N4-5 and 36-40). Labels in X and Y axes are shared by column and rows.

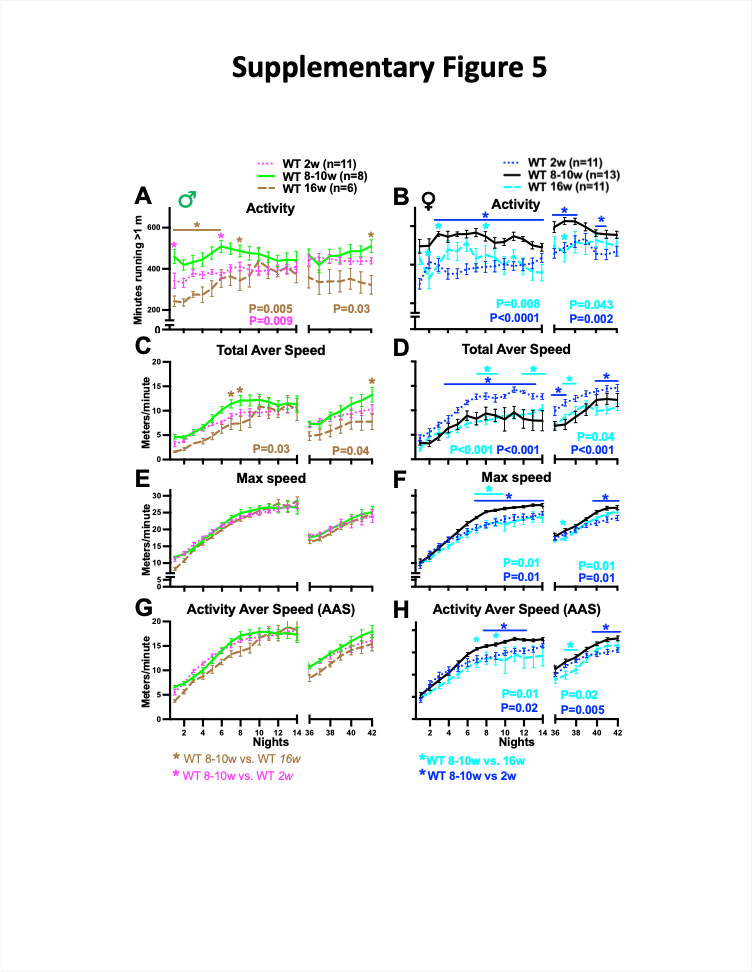

Fig. S5. Age-Driven Changes in CW Performance in WT Male and Females Mice.

Plots for nightly values of CW parameters in WT male (left panels) and female (right panels) mice. The activity levels (A-B), total average speed (C-D; TAS), maximum speed (E-F; MaxSpd), and activity average speed (G-H; AAS) of WT mice subjected to the CW at 2-8w post-tmx (solid lines) were compared with those introduced at 16w post-tmx (dashed lines; brown for males and cyan for females) and at 2w post-tmx (dotted lines; magenta for males and blue for females). Comparisons with statistically significant differences show P values under the plots; color code: 8-10w vs. 16w post-tmx; brown for males and cyan for females, 8-10w vs. 2w post-tmx; magenta for males and blue for females (two-way ANOVA test, with age as source of variation). Significantly lower CW values for individual nights (Bonferroni’s post-hoc, *p<0.05 for all parameters) are shown with the same color code. Labels in the X and Y axes are shared by columns and rows.

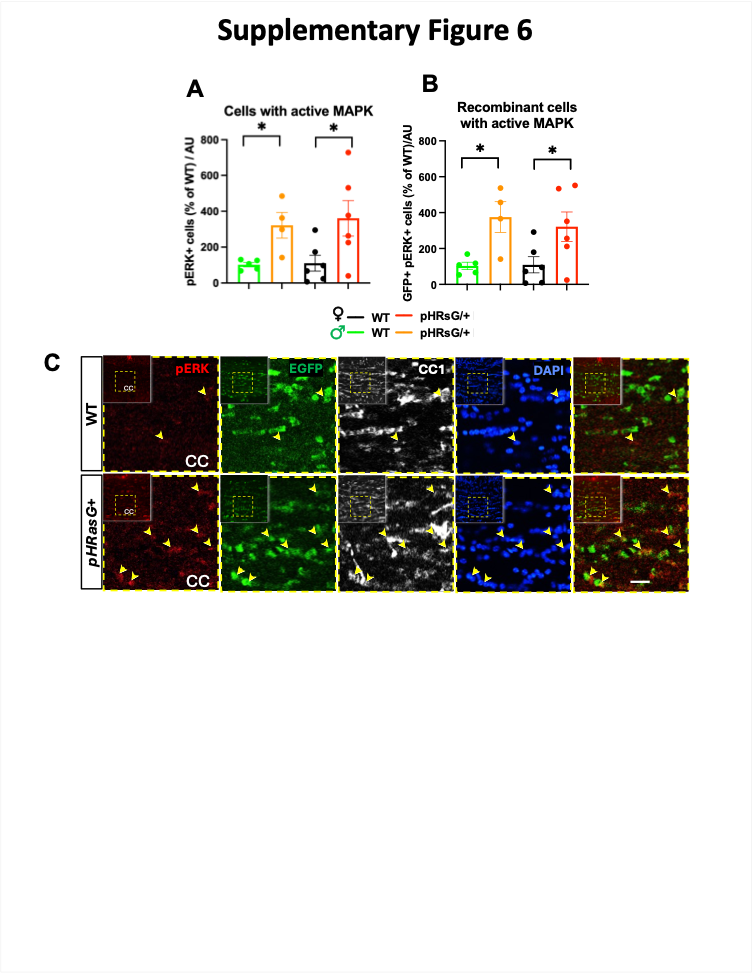

Fig. S6. Functional Impact of OL *HRasG12V* at 2w post-tmx.

The abundance of total (A) and recombinant (B: EGFP+) cells with immunodetected active MAPK (pERK+) in WT and pHRsG/+ females and males is shown (*p< 0.05; unpaired Student’s *t* test). Genotype/sex color code is shown at the bottom. C) Representative immunostainings showing pERK (red), recombinant (EGFP+; green), OLs (CC1+, white), and nuclei (DAPI+; blue) in the corpus callosum (CC) of WT and HRsG/+ mice. Scale bar = 25μm.

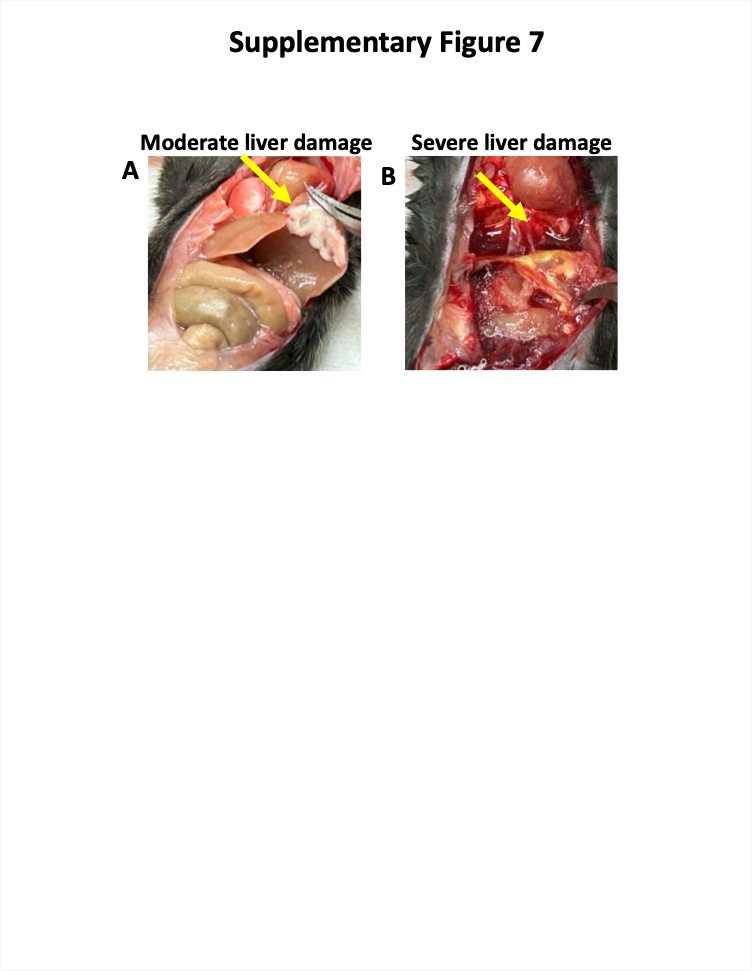

Fig. S7. Liver Damage Following 14-day 1400w Treatment (2mg/kg/day).

The effects of 14-day intraperitoneal treatment with the NOS2 specific inhibitor 1400w in adult pHRsG/+ mice is shown. Moderate (left) and severe (right) liver damage are shown with yellow arrows.

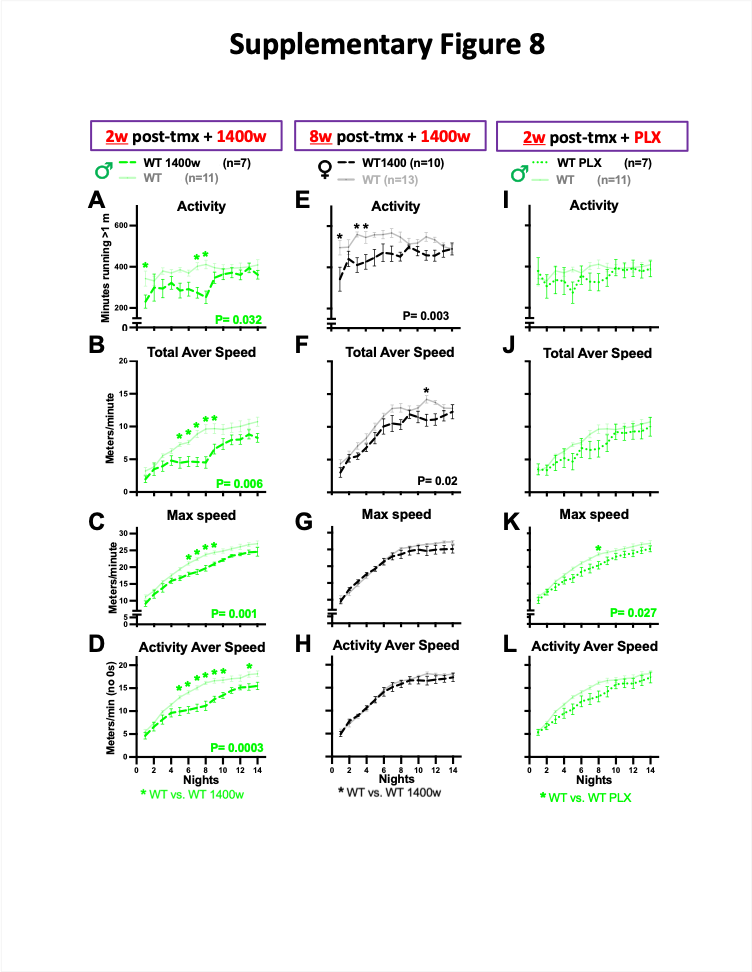

**Fig. S8. Impact of 1400w and PLX3397 on CW Performance in WT Mice.**

Plots for the nightly CW values in WT male (A-D; I-L, green) and female (E-H; black) mice treated with 1400w at 2w (left) and 8w (center) post-tmx, or PLX3397 (PLX) at 2w post-tmx (right). Significantly different P values from comparing 1400w-treated WTs (dashed lines) vs. untreated WTs (solid faded lines) are shown (two-way ANOVA test, treatment as source of variation, A, E; activity levels, B, F; total average speed, C, G; maximum speed, and D, H; activity average speed). I-L) Plots for the nightly CW values in WT males fed with a diet containing PLX for three weeks, as compared with untreated WTs (solid faded lines). The max speed in PLX-treated mice was lower as compared with untreated WTs (K; P=0.027). For all comparisons, Bonferroni’s post-hoc was performed (*p<0.05). The “n” per group is shown in the genotype/sex color code. Labels in X and Y axes are shared in panels of the same column or row.

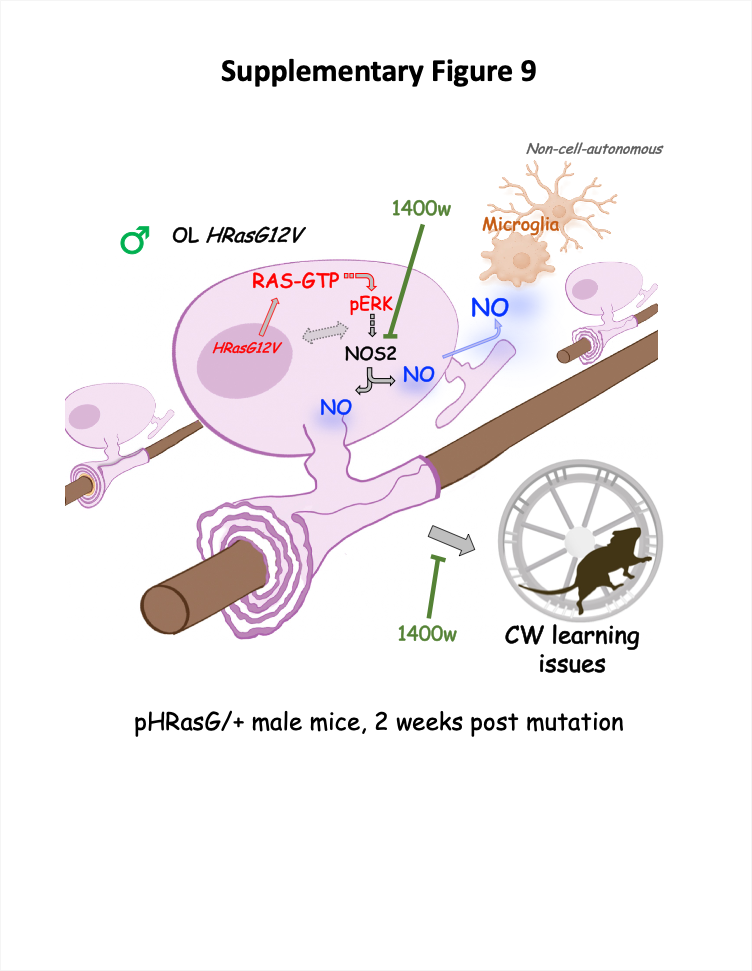

Fig. S9. Working Hypothesis: Impact of *HRasG12V* Mutation in OLs on Molecular Signaling, Cell Physiology, and Fine Motor Skill Learning.

The cartoon illustrates an oligodendrocyte (OL) carrying the *HRasG12V* mutation (male pHRsG/+ mouse) which leads to hyperactive Ras/MAPK pathway (red) and subsequently to increased NOS2 activity. Faded dotted arrow indicates potential transcriptional changes controlling phenotypes. Increased nitric oxide (NO; blue) production acts locally, impacting OL functions including formation of compact myelin (bottom), and diffuses extracellularly affecting neighbor OLs and other cell types (non-cell-autonomous effects on microglia). A key effect mediated by increased NO is the impairment of fine motor skill learning in the complex wheel (CW) test, which can be rescued by inhibition of the NOS2 with the drug 1400w (green).
